# Systems-wide analysis unravels the new roles of CCM signal complex (CSC)

**DOI:** 10.1101/631424

**Authors:** Johnathan Abou-Fadel, Mariana Vasquez, Brian Grajeda, Cameron Ellis, Jun Zhang

**Author notes:** All correspondence: Jun Zhang, Sc.D., Ph.D., Department of Molecular and Translational Medicine (MTM), Texas Tech University Health Science Center El Paso, 5001 El Paso Drive, El Paso, TX 79905.

## Abstract

Cerebral cavernous malformations (CCMs) are characterized by abnormally dilated intracranial capillaries that result in increased susceptibility to stroke. Three genes have been identified as causes of CCMs; KRIT1 (CCM1), MGC4607 (CCM2) and PDCD10 (CCM3); one of them is disrupted in most CCM cases. It was demonstrated that both CCM1 and CCM3 bind to CCM2 to form a CCM signaling complex (CSC) to modulate angiogenesis. In this report, we deployed both RNA-seq and proteomic analysis of perturbed CSC after depletion of one of three CCM genes to generate interactomes for system-wide studies. Our results demonstrated a unique portrait detailing alterations in angiogenesis and vascular integrity. Interestingly, only in-direct overlapped alterations between RNA and protein levels were detected, supporting the existence of multiple layers of regulation in CSC cascades. Notably, this is the first report identifying that both β4 integrin and CAV1 signaling are downstream of CSC, conveying the angiogenic signaling. Our results provide a global view of signal transduction modulated by the CSC, identifies novel regulatory signaling networks and key cellular factors associated with CSC.

## Introduction

Cerebral cavernous malformations (CCMs), abnormally dilated intracranial capillaries, result in focal neurological defects and increased susceptibility to strokes, presenting patients with seizures, headaches and cerebral hemorrhage ^1^. Roughly around 0.5% of the general population are said to have these microvascular malformations, although this number is estimated to be low since many patients with CCMs are asymptomatic ^2^. Familial forms of the disease, accounting for around 55% of patients, results from a predisposition of genetic mutations in three known CCM loci while the etiology of remaining 45% of CCM cases are still unknown ^3-5^. Interestingly, the Hispanic population has a higher percentage (1 in 70) of harboring a potential mutation in one of the three CCM genes compared to non-Hispanic populations (1 in 200) ^6^. CCM induced dilated capillaries, detected in the clinical setting through magnetic resonance imaging (MRI), eventually induce lesions in the central nervous system (CNS), skin and liver. These lesions have characteristic densely packed microvessels and deficient interstitial brain parenchyma ^7^. Treatment for patients suffering from CCM’s ranges from antiepileptic drugs for seizures to surgical excision for those suffering from seizures or hemorrhage ^6^. It has been shown that in addition to cerebral hemorrhagic strokes and seizures, CCM2 variants have been linked to cerebral microbleeds and suggest a link to preclinical Alzheimer’s disease as a result of the CCM2 variant in patients ^8^.

The CCM signaling complex (CSC) is composed of all three CCM genes KRIT1 (CCM1), MGC4607 (CCM2) and PDCD10 (CCM3); in which CCM2 functions as the docking site ^9^. The CSC interacts with other proteins and has affinity for a wide range of ligands; interactions with ligands contribute to cellular processes including cell migration, adhesion and apoptosis ^10-13^. CCM proteins have been shown to alter pathways associated with angiogenesis, the process of new blood vessel development ^14-18^, vascular angiogenesis ^11,19,20^, blood vessel stability ^21^, cell adhesion to extracellular matrix, tube formation and endothelial cell morphogenesis ^16,22^. An association between vascular endothelial growth factor (VEGF) and CCM’s have also been correlated to angiogenesis pathways, creating a more complex understanding of how all these pathways coordinate together ^15,23-25^.

Integrins, cell surface receptors for extracellular matrix (ECM), regulate angiogenesis, along with lymphangiogenesis by promoting events such as pro-angiogenic cell trafficking and endothelial cell (EC) survival and migration ^26^. Integrin signaling is known to be important for angiogenesis, specifically for ECs survival, growth and migration. ^26,27^. Angiogenesis is an essential component of regulating blood brain barrier (BBB) involving integrin ^28^. In fibronectin binding integrins, α5β1 or αvβ3 integrins are known to bind to fibronectin while α5β1 is crucial to regulate levels of αvβ3 in stimulating RhoA-mediated signaling ^29^. In laminin binding integrins, α2β1 are known to enhance vascular endothelial growth factor receptor 1 (VEGFR1) signaling, while α6β1 has been shown to block tube formation and ECs migration ^30^. The β4 integrin, also part of the laminin binding Integrins, are known to mediate hemidesmosome assembly and promote Raf-ERK and Rac-JNK signaling; mice deficient in the β4 integrin C-terminus (cytoplasmic tail) had reduced angiogenic response to VEGF, confirming the essential role of β4 integrin in promoting angiogenesis ^31-33^.

Angiogenesis has also been found to share common signaling pathways involved with inflammation. Inflammation, essential for defending the body from pathogens, has negative effects on tissue including the ability to induce angiogenesis ^34^. It has been shown that regulators of angiogenesis, such as angiopoietin-2 (ANGPT2) upregulates inflammatory responses ^34^. Chemokines and cytokines, which have a large part to play with inflammation, are also important for angiogenesis. One example includes members of the CC chemokine family, such as CCL2, which along with IL-8 mediates localization of endothelial cells to sites of inflammation ^35^. Increased VEGF-A expression by CCL2 leads to induced angiogenesis, which reinforces CCL2’s role in promoting angiogenesis during inflammation ^35^. IL-1 proteins, also involved with inflammatory responses, have been reported to stimulate release of collagenase and prostaglandin from synovial cells, which is known to stimulates expression of VEGF ^36-38^. To our knowledge, this is the first attempt in systems biology, analyzing CCMs deficient models in human brain microvascular endothelial cells (HBMVEC) *in-vitro* and zebrafish lines *in-vivo.* Our results elucidate a very unique and interconnected story demonstrating alterations in various signaling cascades, both in cerebral neovascular angiogenesis as well as developmental angiogenesis with disruption of the CSC signaling cascade. Our goal is to provide a global view of alterations at both the proteome and transcriptome level resulting from deficiencies in CCM expression in both *in-vitro* and *in-vivo* models. Integrative proteomic and genomic analysis of CCM deficiencies in HBMVEC allow us to investigate effects on cerebral microvascular angiogenesis while our zebrafish models allow us to investigate effects on developmental angiogenesis resulting from a disrupted CSC.

## Materials and Methods

### Generation of CCM deficient cells and zebrafish strains

CCM deficient cells were generated by silencing *CCM1, CCM2* and *CCM3* genes respectively in human brain microvascular endothelial cells (HBMVEC) as described ^11,19^ and Ccm1-knockout (Ccm1-KO, *san*) and Ccm2-knockout (Ccm2-KO, *vtn*) zebrafish embryos were produced by crossing the hemizygous parents in our lab as described ^21,22^.

### RNA extraction and RNA-seq for HBMVEC cells and zebrafish embryos

Total RNAs were extracted with TRIZOL reagent (Invitrogen) following the manufacturer’s protocol. For cultured HBMVEC, cell monolayer was rinsed with ice cold PBS once. Cells were lysed directly in a culture dish by adding 1 ml of TRIZOL reagent per flask and scraping with cell scraper. The cell lysate was passed several times through a pipette and vortexed thoroughly. For zebrafish embryos, tissue samples were homogenized in 1 ml of TRIZOL reagent per 50 to 100 mg of tissue, using a glass-Teflon or power homogenizer. The quality (purity and integrity) of each RNA sample was assessed using a Bioanalyzer (Agilent) before RNA-seq. All RNA-seq data were produced using Illumina HiSeq 2000; clean reads for all samples were over 99.5%. 60-80% of reads were mapped to respective reference genomes (Human or zebrafish).

### RNA-seq processing of files to assemble interactomes for HBMVEC and zebrafish embryos

The RNA-seq files for the HBMVEC samples were obtained through paired-end (PE) sequencing with 100 bp reads (2×100) in Illumina HiSeq2000. The data consisted of 8 FASTQ files, 2 PE FASTQ files for each of the four cohorts: KD-CCM1, KD-CCM2, KD-CCM3 and WT (wild type). All cohorts consisted of one sample, with CCM2 acting as a replicate for both CCM1 and CCM3, as CCM2 functions as the docking site for the CSC ^9^. The zebrafish data consisted of 6 FASTQ files, 2 PE FASTQ files for each of the three cohorts: KO-Ccm1 (*san*, Z1), KO-Ccm2 (*vtn*, Z2), and WT (wild type, Zc). Comparisons among the zebrafish samples were made as well (Z1 vs Z2, Z2 vs Zc, and Z1vs Zc). For Zebrafish-Human comparisons, the genes were converted to human homologs and processed the same way as the endothelial cell samples above.

RNA data were cleaned and passed quality control before starting the analysis. After extracting the FASTQ files, the RNA-seq FASTQ files were aligned to the Human genome using HISAT2. This generated two SAM files per cohort that were converted to SAM using SAMtools 1.9. SAMtools quick check was then used to ensure the files were complete. The SAM files were then converted to BAM files and sorted. Cufflinks was used to assemble transcripts for each cohort and cuffmerge to merge all the transcript files. These files were analyzed for differential gene expression with CuffDiff.

After importing the gene exp.diff output file from cuffdiff into Excel, data was filtered for significant 2X and 3X changes in expression. A Python script was created to identify shared differentially expressed genes between cohorts. These were annotated in Excel, and simple set operations were used to find exclusions between the sets. The overlaps were inputted into the Human Reference Protein Interactome Mapping Project database (HURI) and using external links, were sent along with the rest of the genes to Genemania. If there were any identified genes not present in the HURI database, these were copied and manually added to the Genemania network before extracting using Cytoscape. The output from Genemania with query and interactor genes then input into STRING database to obtain the KEGG pathways, GO cellular components, GO molecular functions and GO biological processes.

Identified overlaps were compared between proteomics and RNA-seq data, with KD of either CCM1, CCM2, or CCM3 in HBMVEC and to the zebrafish KO data with KO of Ccm1 (*san*) or Ccm2 (*vtn*). The goal was to identify similarities between cohorts at both the protein and transcription level. Those that were shared with zebrafish were said to be zebrafish validated. The agreements in differential expression were noted through the use of the Python comparison script.

### Protein extraction for HBMVEC cells and zebrafish embryos

HBMVEC cells and zebrafish embryos were lysed using a digital sonifier cell disruptor (Branson model 450 with model 102C converter and double step microtip) in ice cold lysis buffer containing 50 mM Tris-HCl (pH 7.5), 150 mM NaCl, 0.5% NP-40 (Sigma), 50 mM sodium fluoride (Sigma), 1 mM PMSF (Sigma), 1 mM dithiothreitol (Invitrogen) and EDTA-free complete protease inhibitor (Roche). The concentration of protein lysates were measure by Qubit assay (Invitrogen) before analysis.

### Liquid Chromatography-Tandem Mass Spectrometry (LC-MS/MS)

The cell lysates were generated from the four cohorts (KD-CCM1, KD-CCM2, KD-CCM3 and WT) in HBMVEC cells and three cohorts (KO-Ccm1 or *san*, KO-Ccm2 or *vtn*, and WT) in zebrafish embryos. All cohorts consisted of three samples. The cell lysates (20-25 µg of protein) were subjected to trypsin digestion using the FASP method ^39^, following the manufacturer’s protocol (Expedion Inc, San Diego, Ca). Four microliters of each digested sample (100 ng/µl) was loaded onto a 25-cm custom-packed porous silica Aqua C18, 125Å, 5 µm (Phenomenex) column. The porous silica was packed into a New Objective PicoTip Emitter, PF360-100-15-N-5, 15 ± 1 µm and pre-equilibrated with 95% solvent A (100% water, 0.1% formic acid) and 5% solvent B (90% acetonitrile, 10% water, 0.1% formic acid) before injection of the digested peptides, respectively. Liquid Chromatography (LC) separation of the peptides was performed on an Ultimate 3000 Dionex RSLC-nano UHPLC (Thermo Fisher Scientific), equilibrated with 95% solvent A and 5% solvent B (equilibration solution). Samples were loaded onto the column for 10 min, at a constant flow rate of 0.5µL/min, before beginning the elution gradient. Solvent B was then increased from 5% to 35% over 85 min, followed by increase to 95% solvent B over 5 min. The plateau was maintained at 95% solvent B for 9 min, followed by a sharp decrease to 5% solvent B over 1 min. The column was then re-equilibrated with 5% solvent B for 10 min. The total runtime consisted of 120 min. Peptides were analyzed using a Q Exactive Plus Hybrid Quadrupole-Orbitrap Mass Spectrometer (Thermo Fisher Scientific), equipped with a Nanospray Flex Ion Source (Thermo Fischer Scientific). Parameters for the mass spectrometer were as follows: full MS; resolution was set at 70,000 and 17,500, for MS1 and MS2, respectively; AGC target was set at 3e^6^ and 1e^5^ for MS1 and MS2, respectively; Max IT at 50 ms; scan range from 350 to 1600 *m/z* dd-MS^2^; Max IT at 100 ms; isolation window at 3.0.

### Processing of proteomics analysis data to assemble interactomes for perturbed CSC

Proteomic data analysis was performed using the Proteome Discoverer (PD) 2.1.1.21 (Thermo Fisher Scientific), with an estimated false-discovery rate (FDR) of 1%. The Human Database was downloaded in FASTA format on December, 1, 2018, from UniProtKB; http://www.uniprot.org/; 177,661 entries. Common contaminants such as trypsin autolysis fragments, human keratins, and protein lab standards were included in the contaminants database which may be found in the cRAP contaminant database and a couple in house contaminants^40^. The following parameters were used in the PD: HCD MS/MS; fully tryptic peptides only; up to 2 missed cleavages; parent-ion mass tolerance of 10 ppm (monoisotopic); and fragment mass tolerance of 0.6 Da (in Sequest) and 0.02 Da (in PD 2.1.1.21) (monoisotopic). A filter of two-high confidence peptides per protein were applied for identifications. PD dataset was further processed through Scaffold Q+ 4.8.2 (Proteome Software, Portland, OR) to obtain the protein quantification. A protein threshold of 99%, peptide threshold of 95%, and a minimum number of 2 peptides were used for protein validation. The quantitation was performed using emPAI values for comparisons. Statistical analysis was carried out using Student’s *t*-test and a significance level of p<0.05. After files were extracted, they were analyzed through proteome discover (described above) to compare differences between mutant and wild type samples. Identified proteins were pooled together by sample, based on significance of the protein being either under-expressed or over-expressed compared to the control samples. Once these datasheets were available, a Python script was created to identify shared differentially expressed genes between cohorts and used to identify important proteins involved in the CSC, second layer interactors of the CSC, or proteins well-known to have significance in human brain microvascular endothelial cells ^41-47^. Identified proteins were assembled into a separate sheet, which were used to generate the interactomes. Interactomes were made for individual pathways (CSC, CSC second layer and proteins well-known to have significance in human brain microvascular endothelial cells). To simplify the generated interactomes, all identified query proteins were pooled together to generate figures illustrated. In addition, proteins found up-regulated or down-regulated in more than one CCM KD/KO strains were combined to show alterations to these pathways when either of the resulting genes are deficient in expression.

To generate interactomes, identified proteins from above criteria were inputted into HuRI database and using external links, were sent along with the interactors to Genemania (identical to generation of transcriptomes). If there were any identified proteins not present in the HURI database, these were manually added to the Genemania network before extracting using Cytoscape. The output from Genemania with query and interactor proteins was then input into STRING database to obtain the KEGG pathways, GO cellular components, GO molecular functions and GO biological processes identical to the processing of the RNA-seq data.

## Results

### Gene expression profiling by RNA-seq in the deficiency of CCM1

Genomic and interactomic effects of alterations detected through RNA-seq with the knockdown (KD) of CCM1 in HBMVEC are illustrated in Figure 1A. Altered cellular processes related to these genes include various regulation of intrinsic and extrinsic apoptotic signaling pathways, RNA and mRNA processing, RNA and mRNA splicing, spliceosomal tri-snRNP complex assembly, membrane docking, ribonucleoprotein complex assembly, negative regulation of response to stimulus, vesicle docking and regulation of I-κB kinase/NF-κB signaling (Suppl. Table 1A). Affected KEGG pathways include spliceosome and NOD-like receptor signaling pathways (Suppl. Table 1B). Significantly down-regulated genes unique in CCM1 mutant included a neuronal adaptor protein, Amyloid β Precursor Protein Binding Family A Member 2 (APBA2); a group of inflammatory factors: Interleukin 1α (IL1A), Moesin (MSN), C-C Motif Chemokine Ligand 2 (CCL2), BCL2 Related Protein A1 (BCL2A1); and a cellular transporter, Lectin Mannose Binding 1 (LMAN1). Significantly up-regulation of genes, found only in CCM1 deficient cells, were not as prominent as down-regulated genes but included a pre-mRNA splicing factor, Pre-MRNA Processing Factor 8 (PRPF8); two well-known angiogenic factors: Nephroblastoma Overexpressed (NOV), and Kruppel Like Factor 8 (KLF8) and a cellular factor involving autophagy, Dehydrogenase/Reductase X-Linked (DHRSX) (Figure 1A).

**Fig.1.**
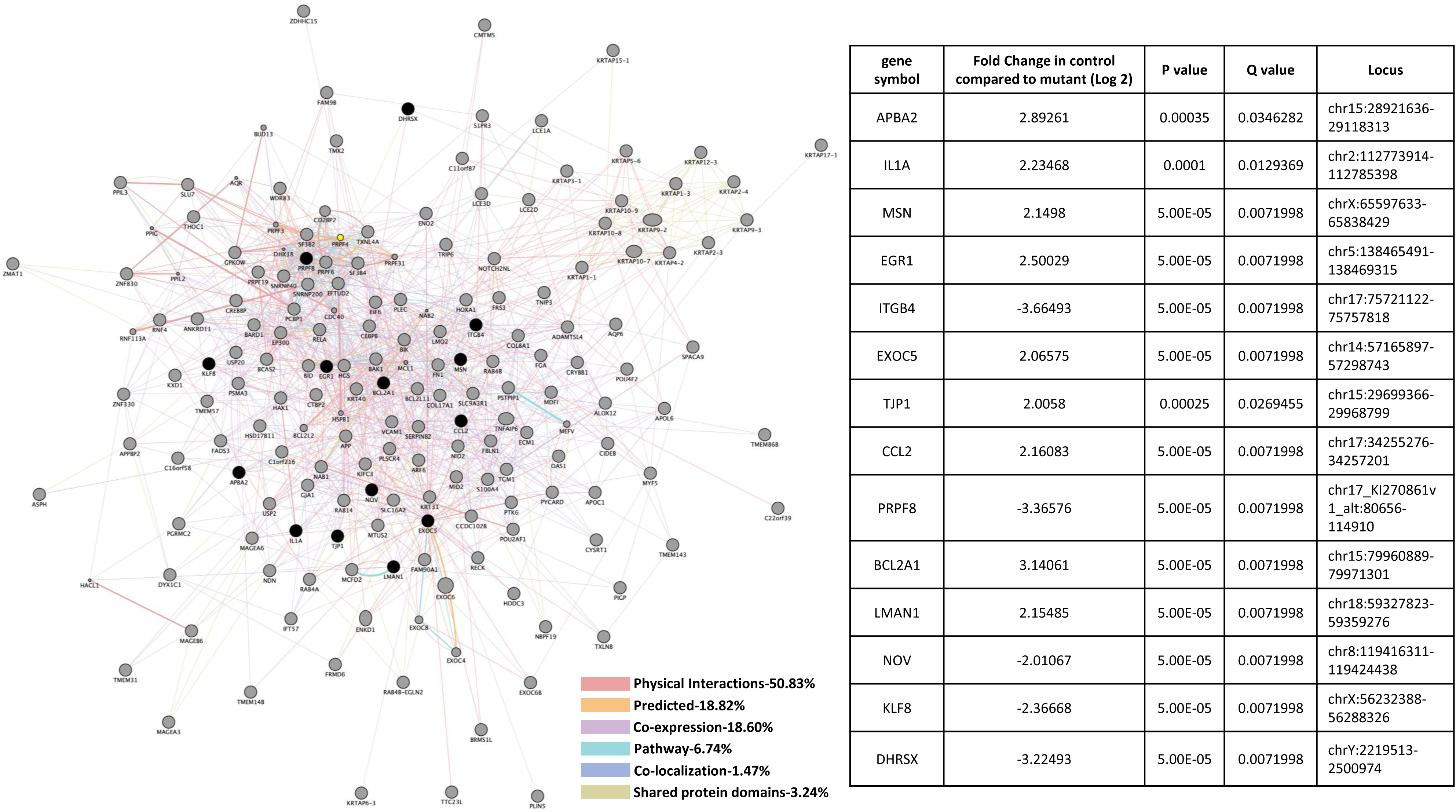
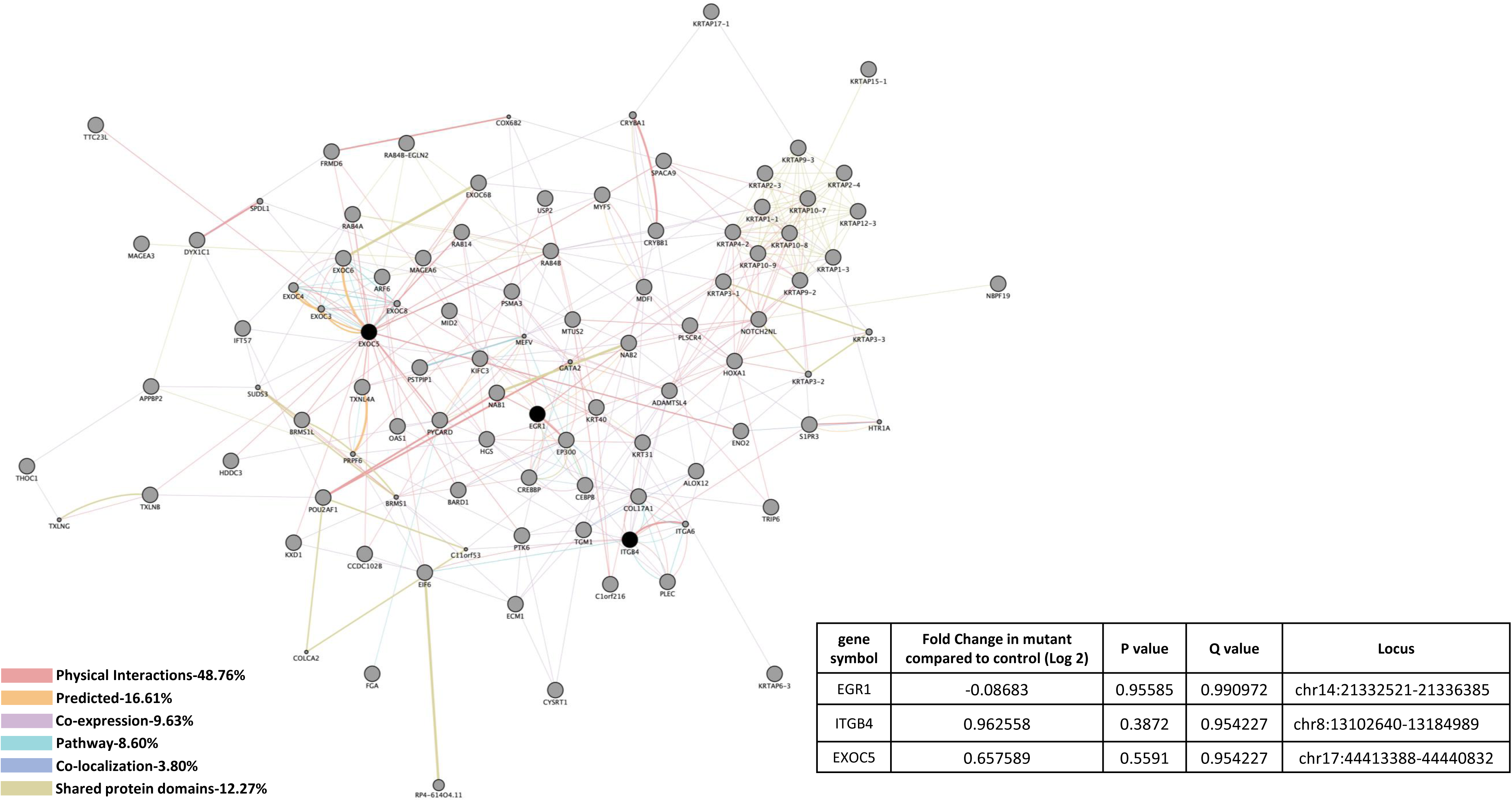
Perturbed transcriptomes in the deficiency of CCM1. A) <u>*CCM1 knockdown human brain microvascular endothelial cells (*HBMVEC).</u> A total of 14 genes were identified with significant expression changes in CCM1 deficient HBMVEC. Data table on right illustrates gene symbol, fold change in control/mutant, P-value and Q-value’s to determine significance and locus identification. B) <u>*CCM1 knockout zebrafish mutant strain (san).*</u> Among the genes identified in HBMVEC, three genes were validated in zebrafish under null-Ccm1 condition. Data table on bottom-right illustrates gene symbol, fold change in mutant/control, P-value and Q-value’s to determine significance and locus identification. For both A and B datasets, RNA-seq results were filtered for significant 2X changes in expression. Interactomes were constructed using cytoscape software equipped with genemania application. Interactomes illustrate affected pathways with both query (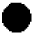) and interactors (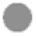). Interaction descriptions with percentages and coordinating line colors are displayed below figures. Significances were determined using Cuffdiff software, depending on whether p > FDR (false discovery rate) after Benjamini-Hochberg correction for multiple-testing and, if Q-value <0.05.

To compare CCM1 interactomes, we also evaluated Ccm1-knockout (KO) zebrafish strain (*san*) and found correlations among 2 genes unique in only CCM1 deficient conditions, either CCM1-KD HBMVEC or CCM1-KO zebrafish (Figure 1A, 1B). Affected cellular processes associated with these two genes included vesicle docking (Suppl. Table 1C). These 2 gene correlations between our models included down-regulation of a transcriptional factor, Early Growth Response 1 (EGR1) in both CCM1-KD HBMVEC cells and Ccm1-KO zebrafish, and down-regulation of a cellular factor in vesicle transport, Exocyst Complex Component 5 (EXOC5) in CCM1-KD HBMVEC cells, but up-regulation in Ccm1-KO zebrafish (Figure 1A, 1B). It must be noted however, that we were unable to obtain significant P-values or Q-values from our RNA-seq analysis with zebrafish data for these two genes (Figure 1B).

### Gene expression profiling by RNA-seq in the deficiency of CCM2

Figure 2A shows pathways affected by alterations in CCM2-KD HBMVEC cells detected by RNA-seq. Affected cellular processes associated with these genes included RNA and mRNA splicing, RNA and mRNA processing, RNA localization and metabolic process, RNA and poly(A) RNA binding, mRNA export from nucleus, mRNA-containing ribonucleoprotein complex export from nucleus, termination of RNA polymerase II transcription, mRNA 3’-end processing, transcription from RNA polymerase II promoter, nitrogen compound transport, ribonucleoprotein complex export from nucleus, ribonucleoprotein complex localization, cell junction organization, cytoplasmic transport, and multiple cellular metabolic processes (Suppl. Table 2A, 2B). Down-regulated genes unique in CCM2 deficiency included an epithelial-to-mesenchymal transition (EMT) marker, Tripartite Motif Containing 44 (TRIM44); and a pre-mRNA splicing factor, U2 Small Nuclear RNA Auxiliary Factor 1 (U2AF1). Interestingly, a cellular regulator of neurotransmitters, Sulfotransferase Family 1A Member 3 (SULT1A3) was the only gene found to be up-regulated unique in CCM2 deficient HBMVEC cells (Figure 2A).

**Fig.2.**
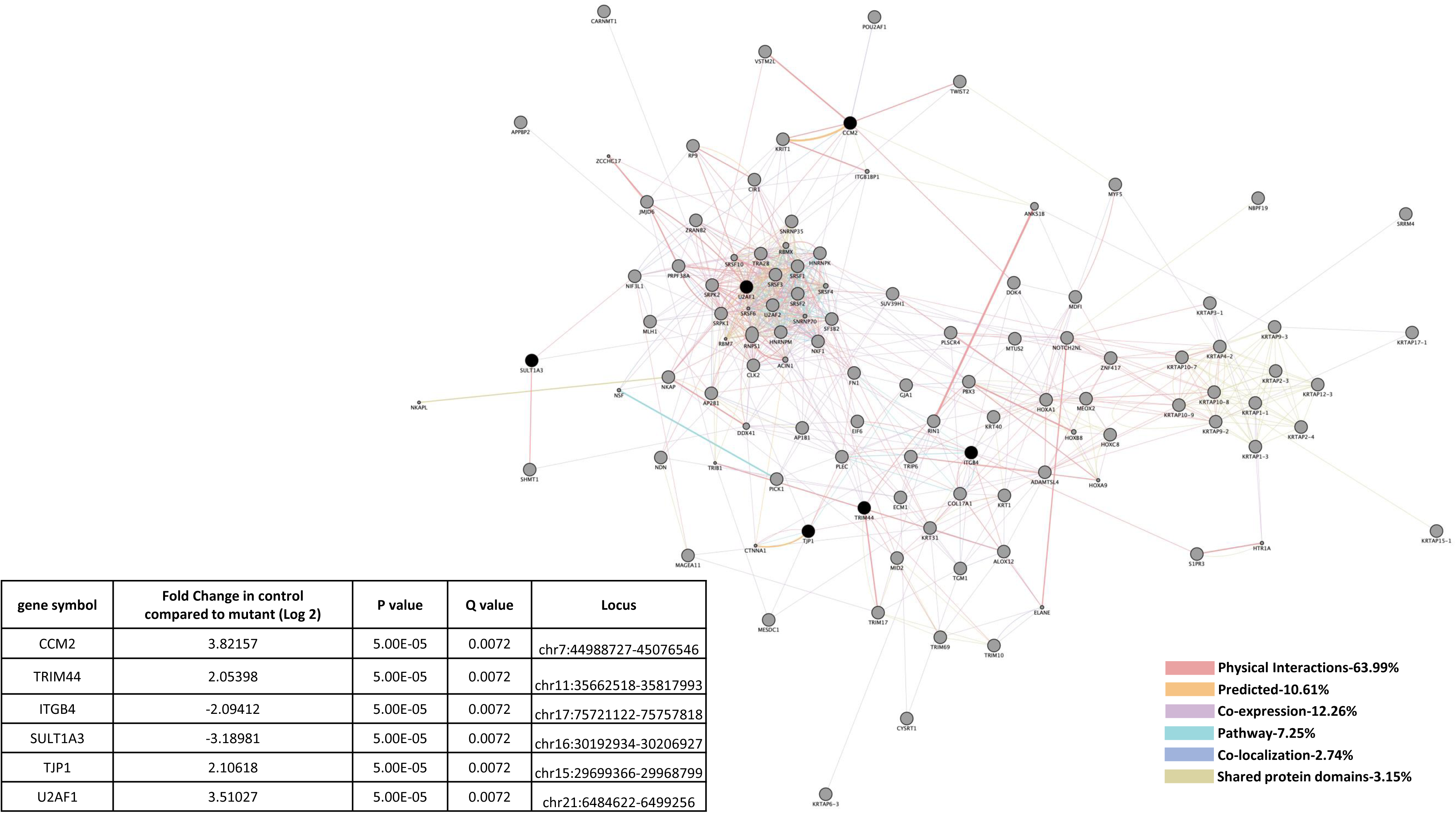
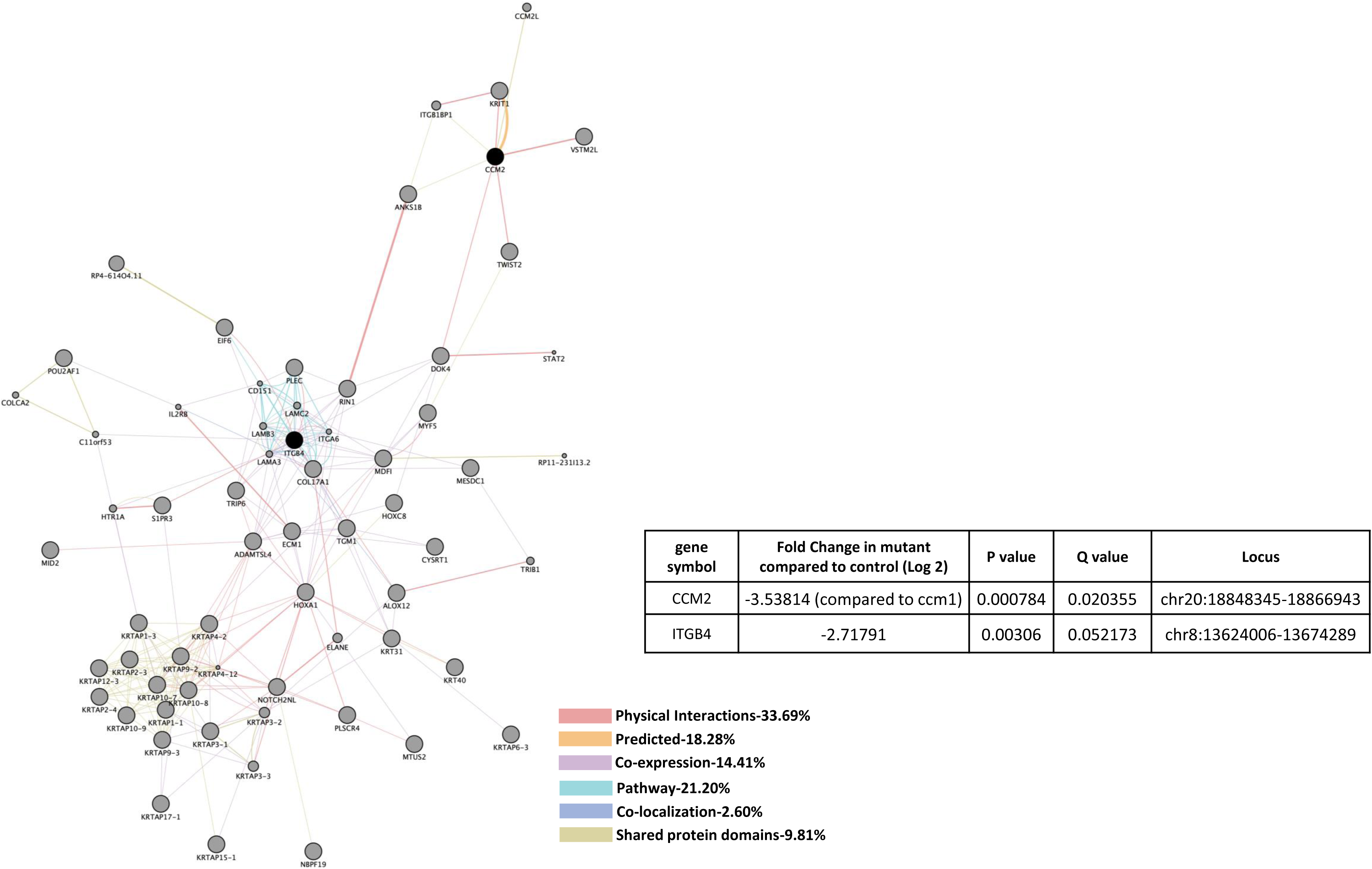
Perturbed transcriptomes in the deficiency of CCM2. A) *<u>CCM2 knockdown human brain microvascular endothelial cells (</u>*<u>HBMVEC).</u> A total of 6 genes were identified with significant expression changes in CCM1 deficient HBMVEC. Data table on left illustrates gene symbol, fold change in control/mutant, P-value and Q-value’s to determine significance and locus identification. B) <u>*CCM2 knockout zebrafish mutant strain (vtn).*</u> Among the genes identified in HBMVEC, two genes were validated in zebrafish under null-Ccm2 condition. Data table on bottom-right illustrates gene symbol, fold change in mutant/control, P-value and Q-value’s to determine significance and locus identification. For both A and B datasets, RNA-seq results were filtered for significant 2X changes in expression. Interactomes were constructed using cytoscape software equipped with genemania application. Interactomes illustrate affected pathways with both query (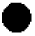) and interactors (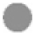). Interaction descriptions with percentages and coordinating line colors are displayed below figures. Significances were determined using Cuffdiff software, depending on whether p > FDR (false discovery rate) after Benjamini-Hochberg correction for multiple-testing, and if Q-value <0.05.

To compare CCM2 interactomes, we also examined expression levels of *CCM2* gene among tissues/cells, and found the expression levels of *CCM2* gene were significantly down in both CCM2-KD HBMVEC cells and Ccm2-KO zebrafish (Figure 2A, 2B), validating the efficiency of our *CCM2* gene silencing (KD) and KO technique, and the accuracy of our RNA-seq data. Affected cellular process associated with CCM2-deficiency target cell junction organization (Suppl. Table 2A, 2B).

### Gene expression profiling by RNA-seq in the deficiency of CCM1 and CCM2

Similarities in altered gene expression in CCM1 and CCM2 deficient HBMVEC cells included down-regulation of Tight Junction Protein 1 (TJP1) (Figure 3A). It should be mentioned that these shared gene alterations were detected separately in individual knockdown experiments, not as a result of a double knockdown strain. Perturbed expression of two EC markers involved in vascular integrity were also observed. First, down-regulation of TJP1 in both CCM1-KD and CCM2-KD HBMVEC cells but was unable to be replicated in *san* or *vtn* zebrafish strains (Figure 3A, 3B). Up-regulation of Integrin β4 (ITGB4), a key component of hemidesmosome, among CCM1-KD and CCM2-KD HBMVEC cells and Ccm1-KO zebrafish (*san*) was observed, but ITGB4 was interestingly down-regulated in Ccm2-KO zebrafish (*vtn*) (Figures 3A, 3B). Affected cellular processes include hemidesmosome assembly, cell junction organization and assembly, cell-substrate junction assembly, cell-matrix adhesion and extracellular matrix organization (Suppl. Table 3A, 3C). Altered KEGG pathways include ECM-receptor interaction, P13K-Akt signaling pathway, and focal adhesion (Suppl. Table 3B, 3D).

**Fig.3.**
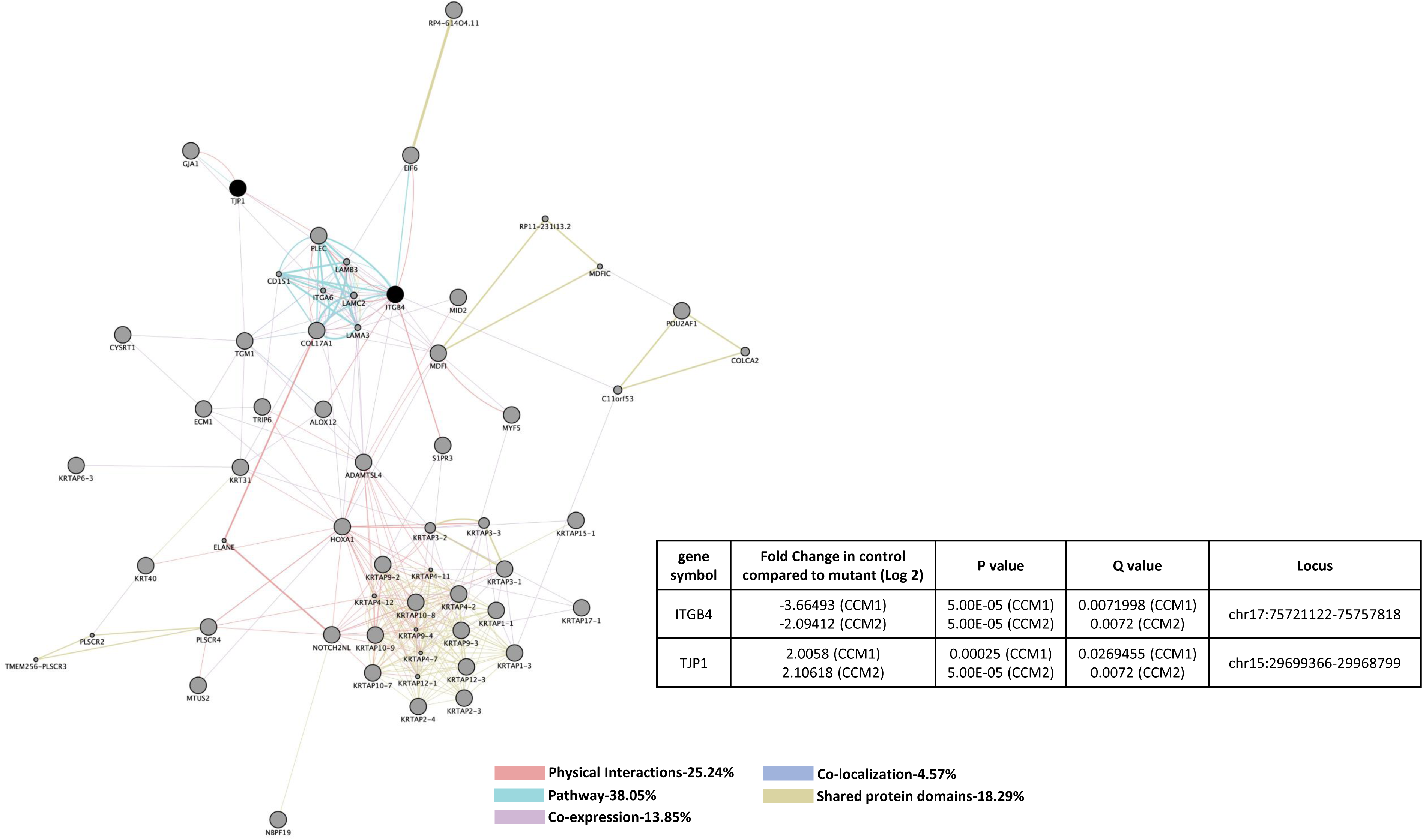
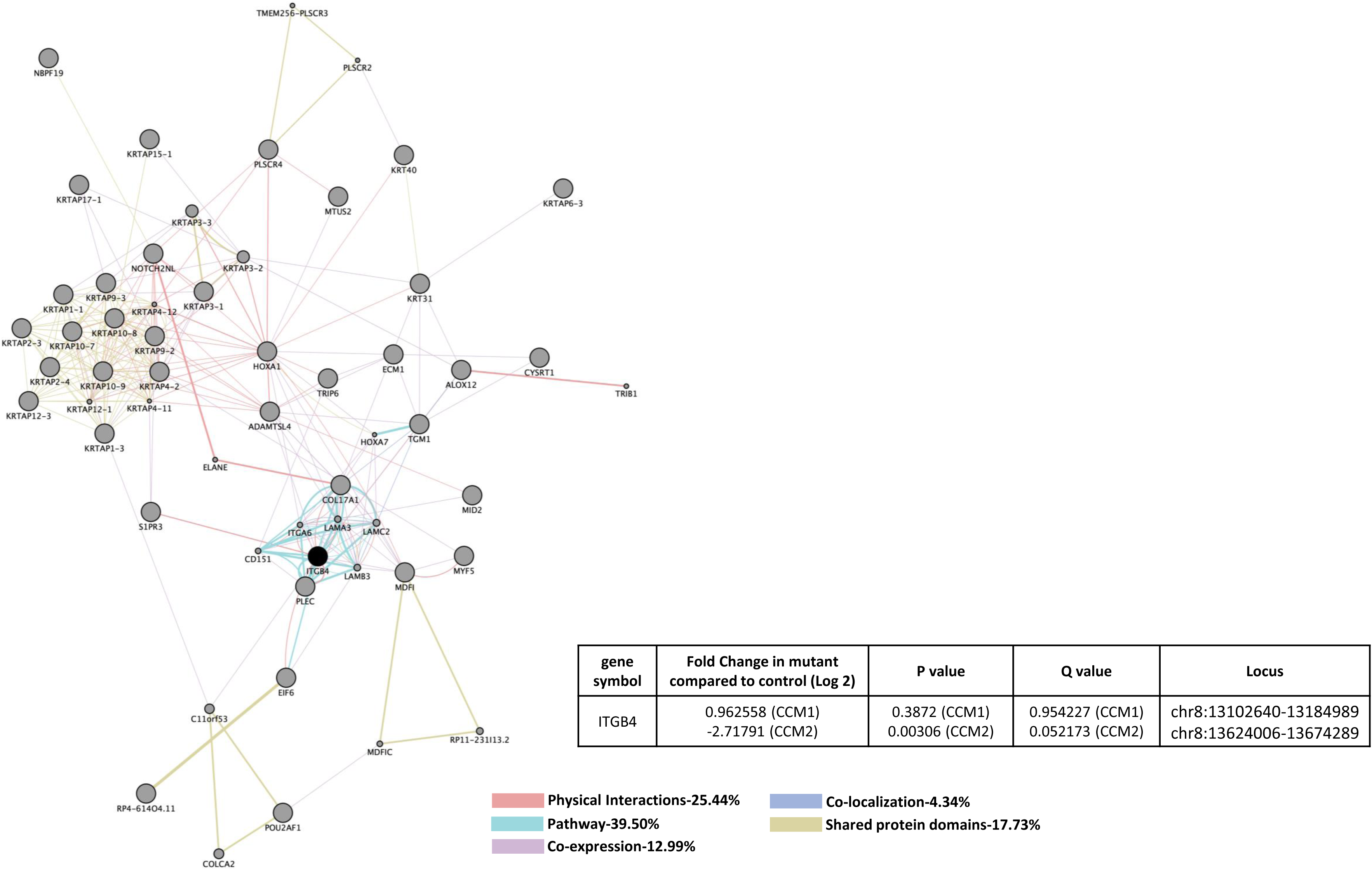
Perturbed transcriptomes shared in the deficiency of CCM1 or CCM2. A) *<u>CCM1 or CCM2 knockdown human brain microvascular endothelial cells (</u>*<u>HBMVEC).</u> Two genes were identified with significant expression changes in both CCM1 and CCM2 deficient HBMVEC. Data table on right illustrates gene symbol, fold change in control/mutant, P-value and Q-value’s to determine significance and locus identification. B) *<u>CCM1 or CCM2 knockout zebrafish mutant strain (san,vtn).</u>* Among the two genes identified in HBMVEC, ITGB4 was validated in zebrafish under null-Ccm1 and Ccm2 conditions. Data table on bottom-right illustrates gene symbol, fold change in mutant/control, P-value and Q-value’s to determine significance and locus identification. For both A and B datasets, RNA-seq results were filtered for significant 2X changes in expression. Interactomes were constructed using cytoscape software equipped with genemania application. Interactomes illustrate affected pathways with both query (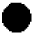) and interactors (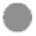). Interaction descriptions with percentages and coordinating line colors are displayed below figures. Significances were determined using Cuffdiff software, depending on whether p > FDR (false discovery rate) after Benjamini-Hochberg correction for multiple-testing, and if Q-value <0.05.

### Gene expression profiling by RNA-seq in the deficiency of CCM3

Heme oxygenase (HMOX2) was the only down-regulated gene detected in CCM3-KD HBMVEC cells (Figure 4). With its highest expression in the cerebrovasculature in the brain ^48,49^, HMOX2 has a role in neovascularization and heme oxygenase (decyclizing) activity (Suppl. Table 4).

**Fig4.**
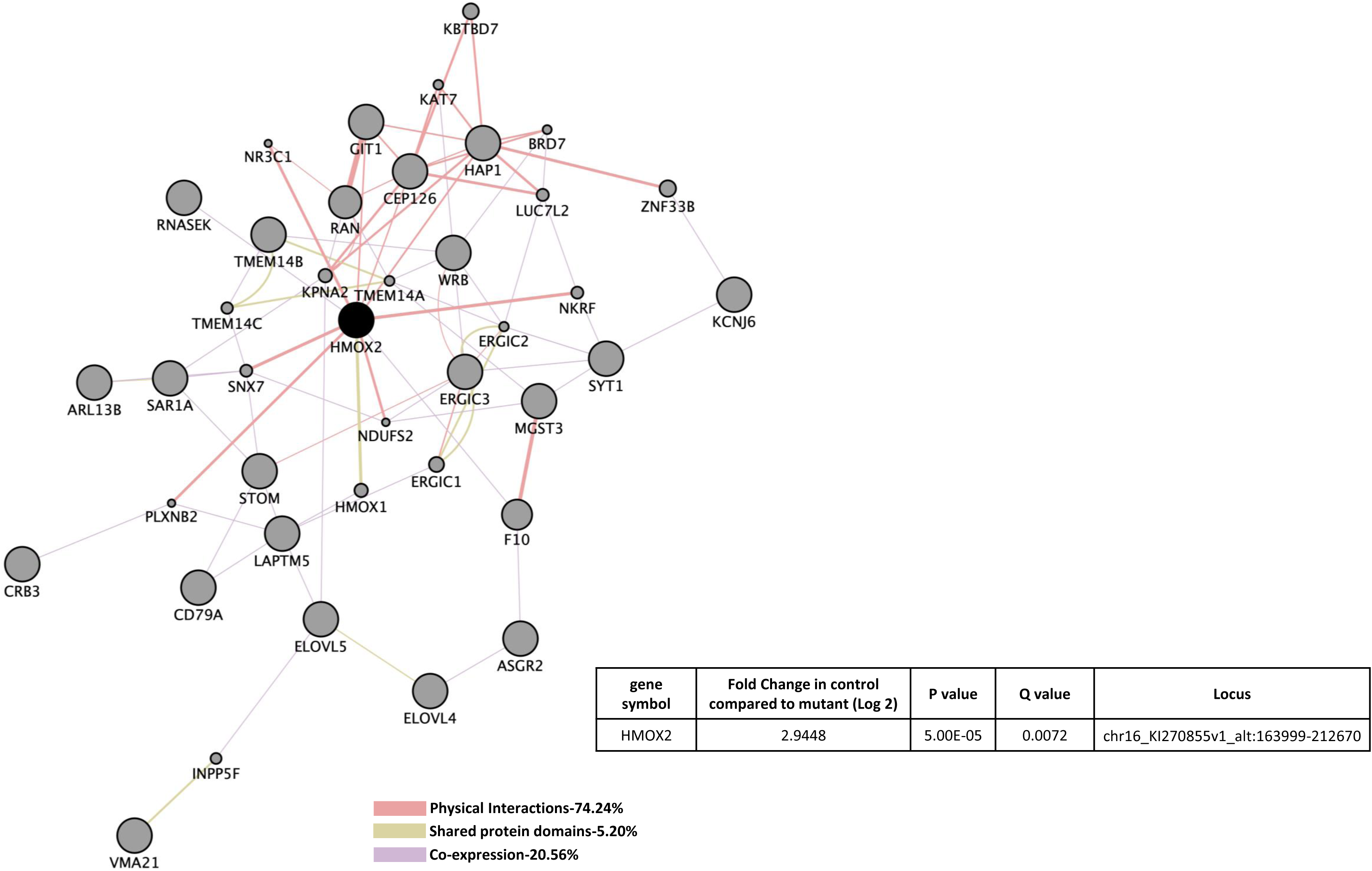
Perturbed transcriptomes in the deficiency CCM3 in human brain microvascular endothelial cells (HBMVEC). HMOX2 was the only identified gene with significant expression changes in CCM3 deficient HBMVEC. Data table on right illustrates gene symbol, fold change in control/mutant, P-value and Q-value’s to determine significance and locus identification. RNA-seq results were filtered for significant 2X changes in expression. Interactome was constructed using cytoscape software equipped with genemania application. Interactome illustrates affected pathways with both query (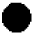) and interactors (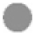). Interaction descriptions with percentages and coordinating line colors are displayed below figure. Significances were determined using Cuffdiff software, depending on whether p > FDR (false discovery rate) after Benjamini-Hochberg correction for multiple-testing, and if Q-value <0.05.

### Perturbed protein interactome pathways in the deficiency of CCM1

Proteins altered that were unique in CCM1-KD HBMVEC cells included down-regulation of Calpain-2 (CAPN2), endothelial cell specific chemotaxis regulator (ECSCR), Glyceraldehyde-3-Phosphate Dehydrogenase (GAPDH) and Cathepsin B (CTSB) (Figure 5A). At the protein level, the majority of altered cellular processes due to CCM1 deficiency lay in the area of vascular angiogenesis, including angiogenesis, blood vessel development and morphogenesis, circulatory system development, tube development and morphogenesis, cellular developmental process, cell differentiation, chemical response, chordate embryonic development, neuron apoptotic process and microtubule cytoskeleton organization (Suppl. Table 5A). Altered molecular functions of interest included cytoskeletal protein binding, ion binding and calcium ion binding (Suppl. Table 5B). Affected KEGG pathways included cellular senescence, Alzheimer’s disease, focal adhesion, apoptosis and HIF-1 signaling pathway (Suppl. Table 5C).

**Fig.5.**
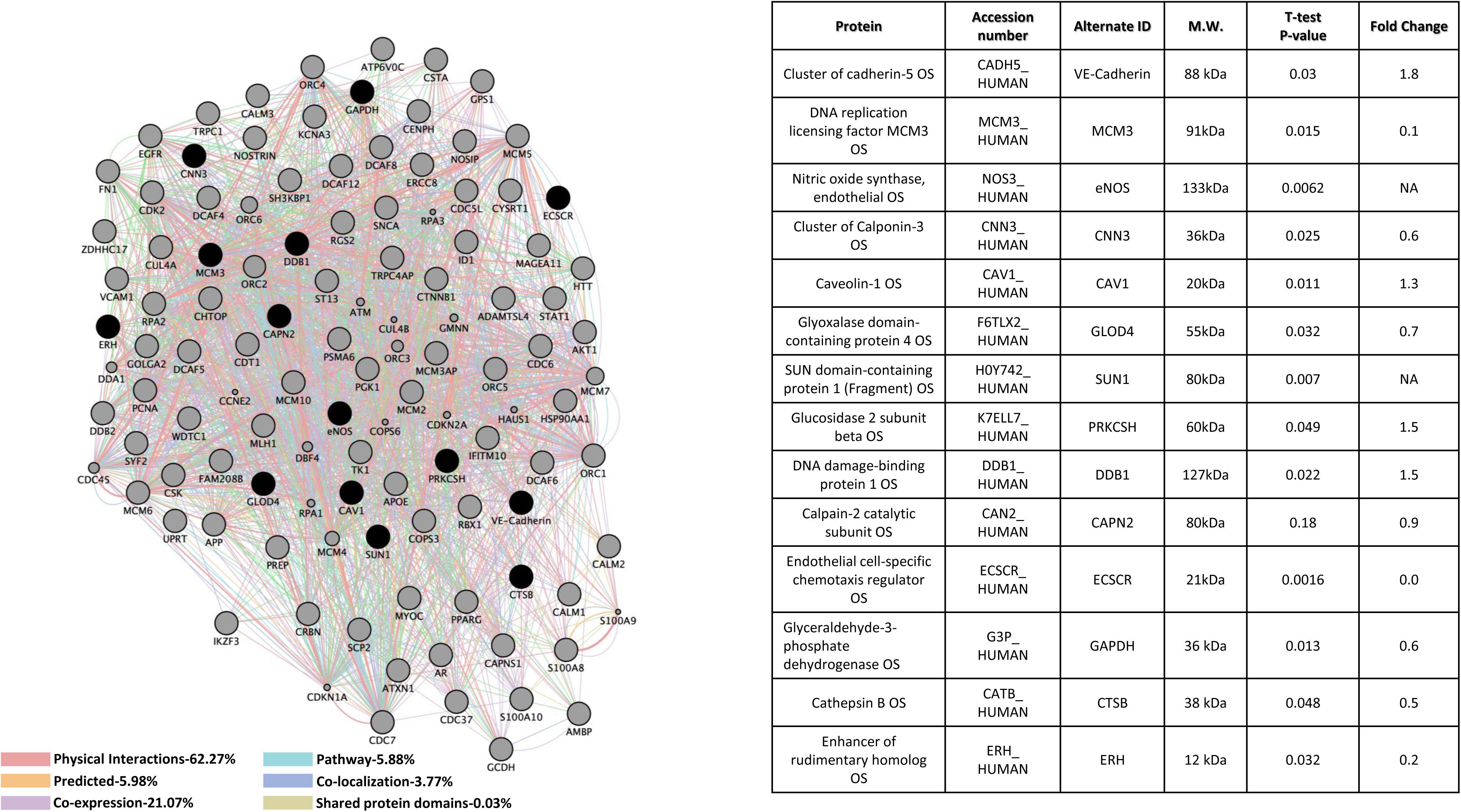
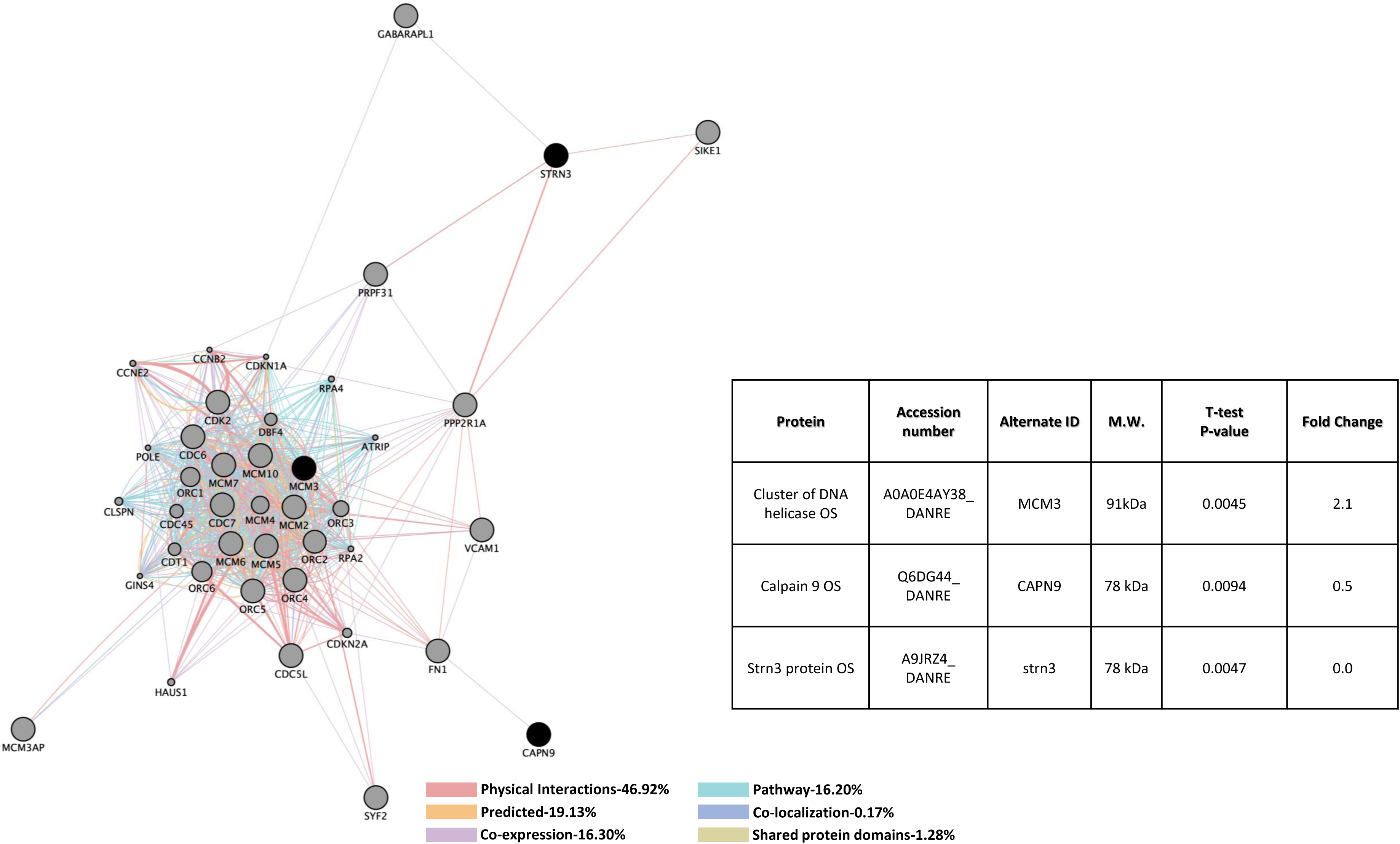
Perturbed protein interactome pathways in the deficiency of CCM1. A). <u>*CCM1 knockdown human brain microvascular endothelial cells (HBMVEC)*</u>. Fourteen proteins were identified with significant expression changes in CCM1 deficient HBMVEC. Table on the right illustrates details of query proteins identified in the samples through mass spectrometry with accession number, alternate ID, Molecular Weight (M.W.), p-values and fold changes in mutant/controls. NA= not found in WT samples. B). <u>*CCM1 knockout zebrafish mutant strain (san)*</u>. Three proteins were identified with significant expression changes in CCM1 deficient zebrafish (*san*). Table on the right illustrates details of query proteins identified in the samples through mass spectrometry with accession number, alternate ID, molecular weight (M.W.), p-values and fold changes in mutant/controls. For both A and B datasets, interactomes were built using proteomic data and constructed using cytoscape software equipped with genemania application. Interactomes illustrate affected pathways with both query (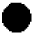) and interactors (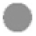). Interaction descriptions with percentages and coordinating line colors are displayed below figures. All samples were run in triplicates. Significances were determined using student’s t-test with p values <0.05.

To compare CCM1 interactomes, we also evaluated Ccm1-KO zebrafish strain (*san*) which gave less interactome data than with CCM1-KD HBMVEC (Figure 5B). Alterations in genes unique in Ccm1-KO zebrafish involved multiple disrupted metabolic processes as well as negative regulation of cellular process (Suppl. Table 5D). Molecular functions potentially affected included binding mechanisms as well as hydrolase activity and catalytic activity (Suppl. Table 5E). Similar to down-regulated Calpain-2 (CAPN2), found in CCM1-KD HBMVEC (Figure 5A), Calpain-9 (CAPN9) was found to be down-regulated in CCM1-KD zebrafish (Figure 5B).

### Perturbed protein interactome pathways in the deficiency of CCM2

In CCM2 deficient HBMVEC, five unique affected proteins were found, including up-regulation of three proteins controlling cellular protein stability: Splicing Factor Proline And Glutamine Rich (SFPQ), Heat Shock Protein Family A Member 4 (HSPA4), Heat Shock Protein Family B (Small) Member 1 (HSPB1) and one blood coagulating factor, Trans aldolase 1 (TALDO1); down-regulation of an RNA-binding protein, Proliferation-Associated 2G4 (PA2G4) was also observed (Figure 6A). Cellular processes potentially affected in the deficiency of CCM2 included cell cycle process, cellular component organization or biogenesis, response to organic substance, various metabolic processes, regulation of oxidative stress-induced intrinsic apoptotic signaling pathway and neutrophil degranulation (Suppl. Table 6A). Molecular functions included transferase activity, transferring aldehyde or ketonic groups and various binding pathways (Suppl. Table 6B). Interestingly, KEGG pathways affected in CCM2 deficient HBMVEC included biosynthesis of amino acids, carbon metabolism, pentose phosphate pathway, antigen processing and presentation and VEGF signaling pathway (Suppl. Table 6C).

**Fig.6.**
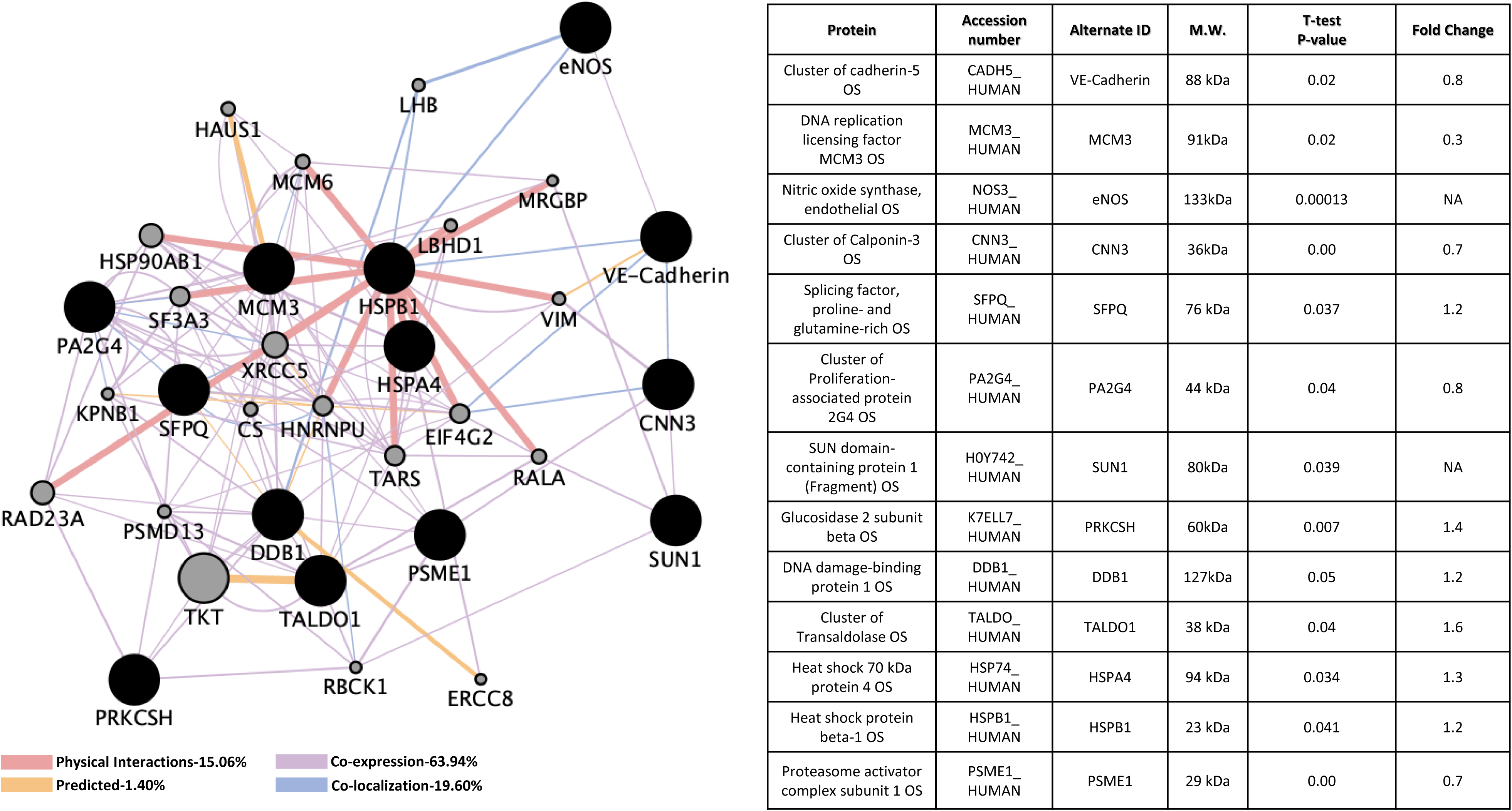
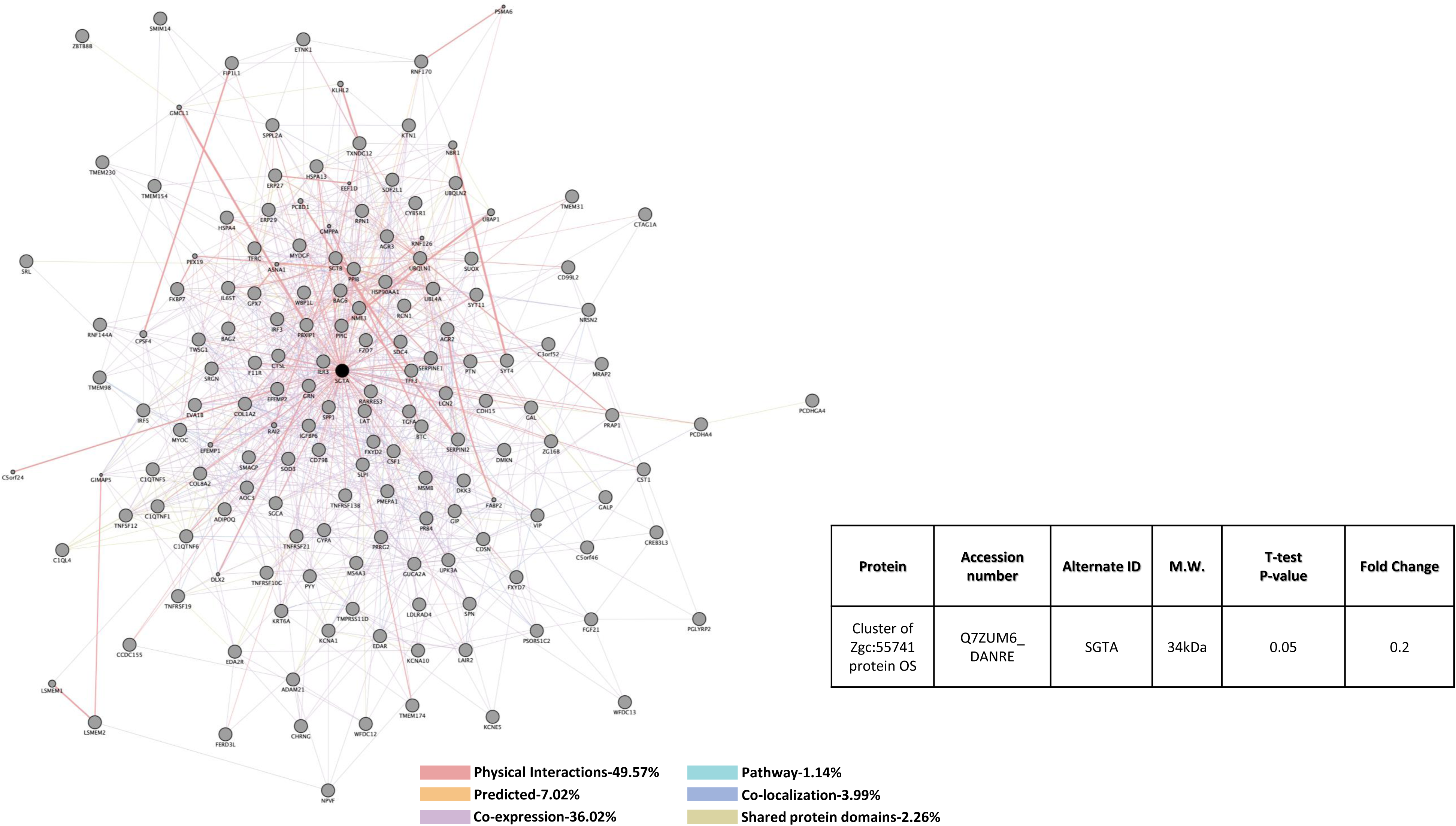
Perturbed protein interactome pathways in the deficiency of CCM2. A). <u>*CCM2 knockdown human brain microvascular endothelial cells (HBMVEC)*</u>. Thirteen proteins were identified with significant expression changes in CCM2 deficient HBMVEC. Table on the right illustrates details of query proteins identified in the samples through mass spectrometry with accession number, alternate ID, Molecular Weight (M.W.), p-values and fold changes in mutant/controls. NA= not found in WT samples. B). <u>*CCM2 knockout zebrafish mutant strain (vtn)*</u>. SGTA was identified with significant expression changes in CCM2 deficient zebrafish (*vtn*). Table on the right illustrates details of query proteins identified in the samples through mass spectrometry with accession number, alternate ID, molecular weight (M.W.), p-values and fold changes in mutant/controls. For both A and B datasets, interactomes were built using proteomic data and constructed using cytoscape software equipped with genemania application. Interactomes illustrate affected pathways with both query (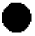) and interactors (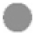). Interaction descriptions with percentages and coordinating line colors are displayed below figures. All samples were run in triplicates. Significances were determined using student’s t-test with p values <0.05.

Only one down-regulated protein, Small Glutamine Rich Tetratricopeptide Repeat Containing Alpha (SGTA), in Ccm2-null zebrafish (*vtn*) was identified (Figure 6B), which was not observed with CCM2 deficient HBMVEC (Figure 6A). Cellular processes associated with SGTA included cellular regulations in response to stimulus, in responses to stress; in responses to metabolic and catabolic processes and ERAD pathway regulation (Suppl. Table 6D). Molecular functions of SGTA included protein binding and are mainly associated with protein stability and ER-associated autophagy processes (Suppl. Table 6E).

### Perturbed protein interactome pathways in the deficiency of CCM1 and CCM2

There was an overlapped group of proteins that were perturbed in deficiencies in either CCM1 or CCM2 expression in HBMVEC (Figure 7A). Among these shared group of altered genes, Cadherin 5 (CDH5, VE-Cadherin) is up-regulated in CCM1-KD but slightly down-regulated in CCM2-KD. VE-Cadherin, a key player in endothelial adherens junction assembly and maintenance ^50^, is involved in BBB integrity, ERK signaling and immune cell transmigration through VCAM-1/CD106 signaling pathways ^51-53^. Nitric oxide synthase 3 (NOS3, eNOS) is up-regulated in both CCM1-KD and CCM2-KD HBMVEC cells. eNOS participates in VEGF induced angiogenesis ^54^, while altered eNOS expression levels are associated with strokes ^55-59^. Both Calponin-3 (CNN3) that is highly expressed in brain and Minichromosome Maintenance Complex Component 3 (MCM3) that is involved in DNA replication ^60,61^ are down regulated in both CCM1-KD and CCM2-KD HBMVEC cells. Cellular processes involving these 4 proteins include various cell cycle pathways, numerous metabolic pathways, VEGFR signaling pathway, negative regulation of ion transport, regulation of cell proliferation and cell activation, second-messenger-mediated signaling, blood circulation and coagulation, RTK signaling pathway, *in-utero* embryonic development, negative regulation of platelet activation, regulation of heart contraction, blood vessel development and various DNA replication pathways (Suppl. Table 7A). Molecular functions of these proteins mainly included various binding pathways, DNA helicase activity and nucleoside-triphosphatase activity (Suppl. Table 7B). Potentially affected KEGG pathways included cell cycle, DNA replication, PI3K-Akt signaling pathway, estrogen signaling pathway and HIF-1 signaling pathway (Suppl. Table 7C).

**Fig. 7.**
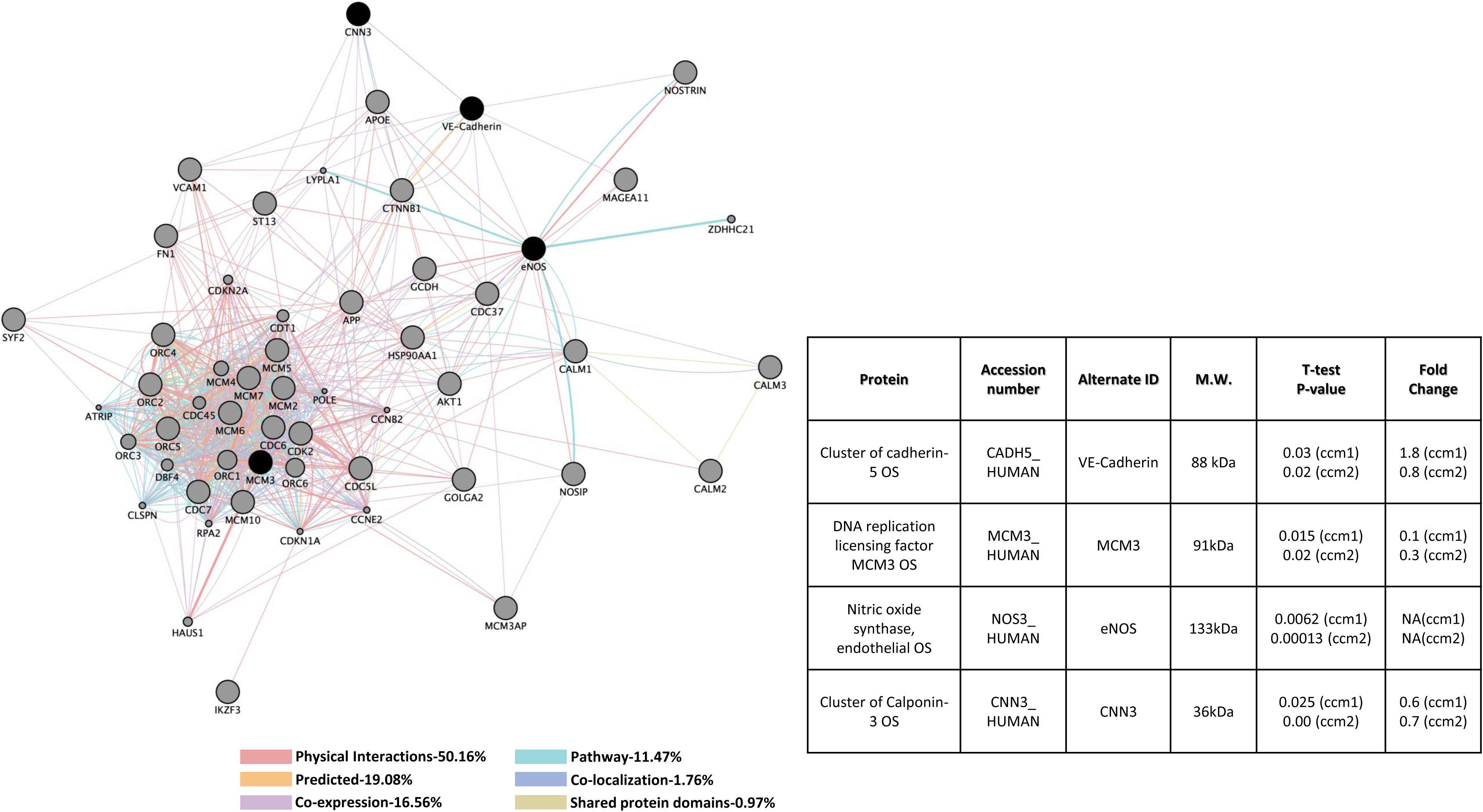
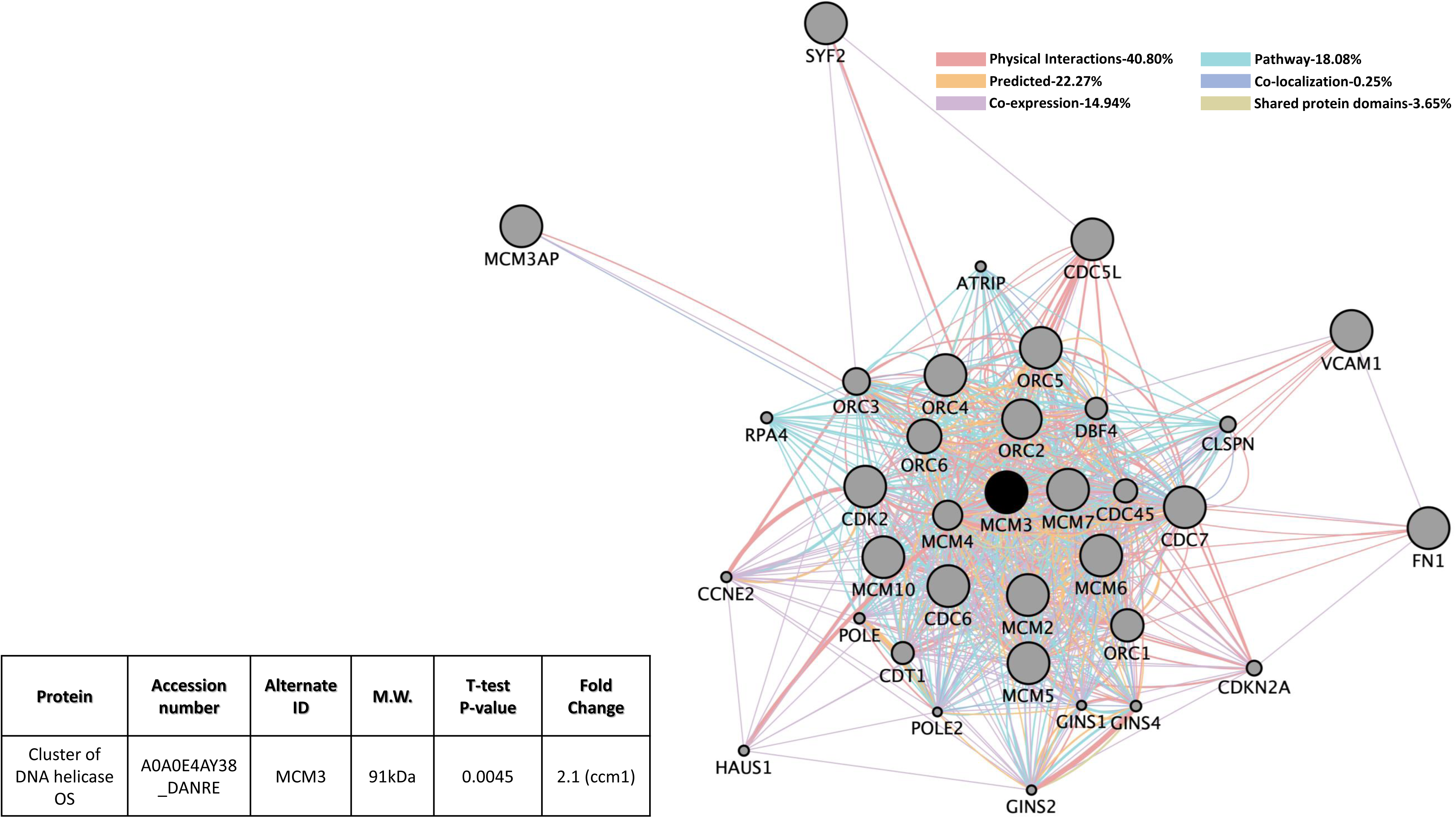
Perturbed protein interactome pathways in the deficiency of CCM1 or CCM2. A). <u>*CCM1 or CCM2*</u> *<u>knockdown human brain microvascular endothelial cells (HBMVEC)</u>*. Four proteins were identified with significant expression changes in CCM1 or CCM2 deficient HBMVEC. Table on the right illustrates details of query proteins identified in the samples through mass spectrometry with accession number, alternate ID, Molecular Weight (M.W.), p-values and fold changes in mutant/controls. NA= not found in WT samples. B). <u>*CCM1 knockout zebrafish mutant strain (san)*</u>. MCM3 was identified with significant expression changes in only CCM1 deficient zebrafish (*san*). Table on the left illustrates details of query proteins identified in the samples through mass spectrometry with accession number, alternate ID, molecular weight (M.W.), p-values and fold changes in mutant/controls. For both A and B datasets, interactomes were built using proteomic data and constructed using cytoscape software equipped with genemania application. Interactomes illustrate affected pathways with both query (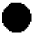) and interactors (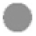). Interaction descriptions with percentages and coordinating line colors are displayed below figures. All samples were run in triplicates. Significances were determined using student’s t-test with p values <0.05.

MCM3 is the only gene found to have shared down-regulation in protein level in CCM1 and CCM2 mutants in HBMVEC. MCM3 protein level was also altered in Ccm1-KO zebrafish (Figures 7A and 7B). This alteration, however, was only demonstrated for Ccm1-KO zebrafish but not Ccm2-KO zebrafish (Figure 7B). It must also be noted that MCM3, in CCM1-KO zebrafish, was up-regulated but was down-regulated in both CCM1-KD and CCM2-KD HBMVEC cells (compare Figures 7A and 7B), perhaps suggesting a different level of feed-back regulation for this protein between cerebral vascular angiogenesis and developmental angiogenesis. Cellular processes in CCM1-KD and CCM2-KD HBMVEC cells validated in Ccm1-KO zebrafish included metabolic processes, DNA replication processes and various cell cycle processes (Suppl. Table 7D). Zebrafish conferred molecular functions mainly included binding pathways and DNA helicase activity (Suppl. Table 7E). Validated KEGG pathways included cell cycle and DNA replication (Suppl. Table 7F).

### Perturbed protein interactome pathways in the deficiency of CCM3

Down-regulated expression of two interesting proteins, unique in CCM3-deficinet HBMVEC cells, Serine/Threonine Kinase 24 (STK24) and Procollagen-Lysine, 2-Oxoglutarate 5-Dioxygenase 2 (PLOD2), were observed (Figure. 8). STK24 is among the first group of proteins identified to bind to CCM3 protein ^62,63^, which further validates our current data. Metabolic processes, various regulations of kinase activity, various responses to stress, regulation of mitotic cell cycle, regulation of MAPK cascade, response to stimulus, regulation of transferase activity, intracellular signal transduction processes, activation of MAPKKK activity and various apoptotic signaling pathways are associated with these two genes (Suppl. Table 8A). Molecular functions affected included MAPKKKK activity (Suppl. Table 8B).

**Fig.8.**
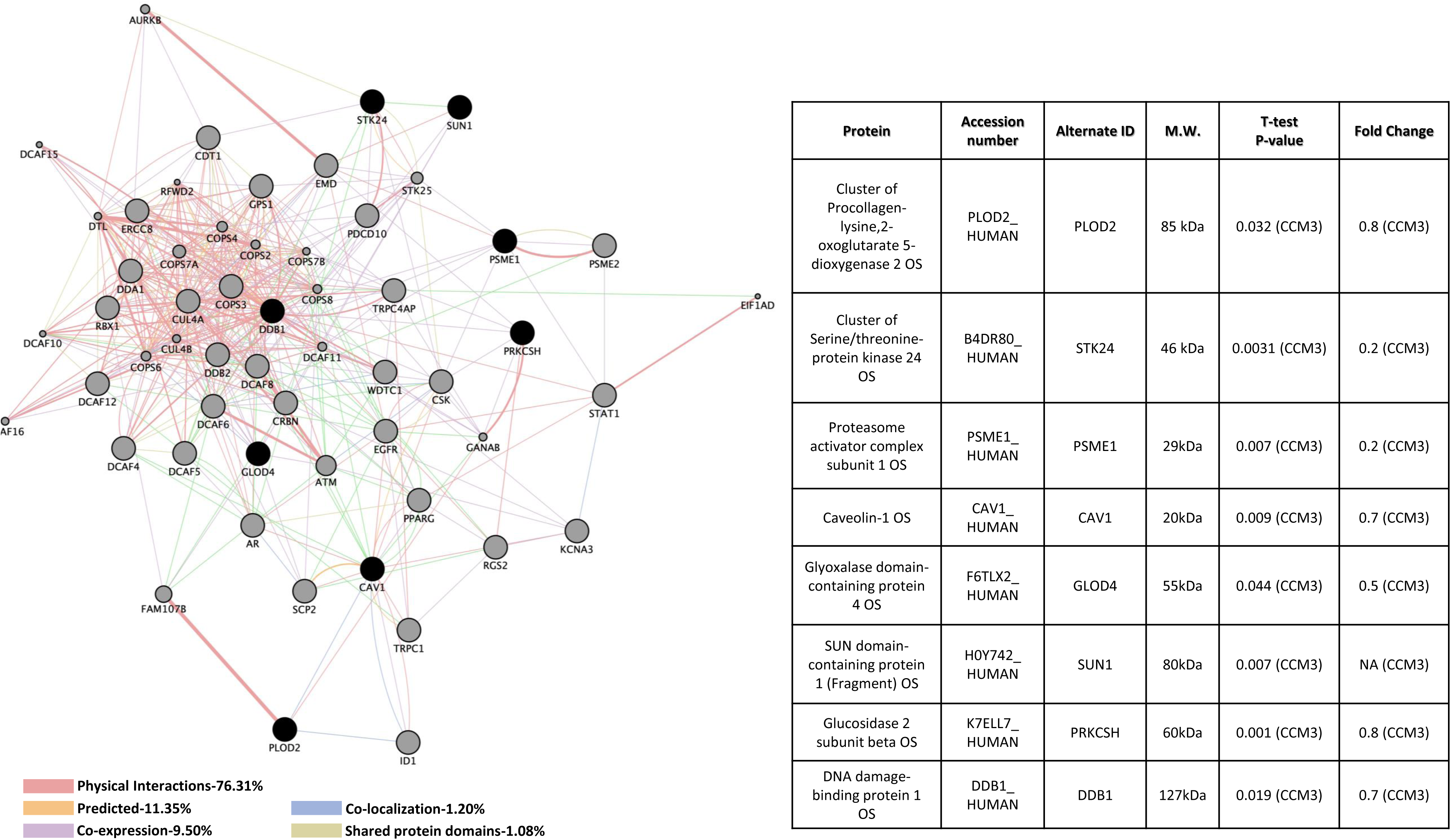
Perturbed protein interactome pathways in the deficiency of CCM3 in human brain microvascular endothelial cells (HBMVEC). Eight proteins were identified with significant expression changes in CCM3 deficient HBMVEC. Table on the right illustrates details of query proteins identified in the samples through mass spectrometry with accession number, alternate ID, Molecular Weight (M.W.), p-values and fold changes in mutant/controls. NA= not found in WT samples. Interactome was built using proteomic data and constructed using cytoscape software equipped with genemania application. Interactome illustrates affected pathways with both query (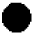) and interactors (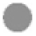). Interaction descriptions with percentages and coordinating line colors are displayed below figure. All samples were run in triplicates. Significances were determined using student’s t-test with p values <0.05.

### Perturbed protein interactome pathways in the deficiency of CCM1 and CCM3

Up-regulation of the scaffolding protein, Caveolin-1 (CAV1), was observed for CCM1 mutants, while down-regulation in CCM3 mutants was observed (Figure 9), suggesting a potentially intricate feed-back regulation within the CSC. Interestingly, down-regulation of Glyoxalase Domain Containing 4 (GLOD4) was shared in both CCM1 and CCM3 mutants (Figure 9). Cellular processes associated with these two genes were vast, but included neurogenesis, nervous system development, cellular response to hormone stimulus, negative regulation of signal transduction, circulatory system development, regulation of cell proliferation, apoptotic processes, membrane biogenesis, regulation of MAPK cascade, caveola assembly, blood vessel development and morphogenesis and vasculature development (Suppl. Table 9A). Molecular functions potentially affected mainly included binding pathways (Suppl. Table 9B). The main KEGG pathway of interest affected for these mutants included focal adhesion pathway (Suppl. Table 9C).

**Figure 9:**
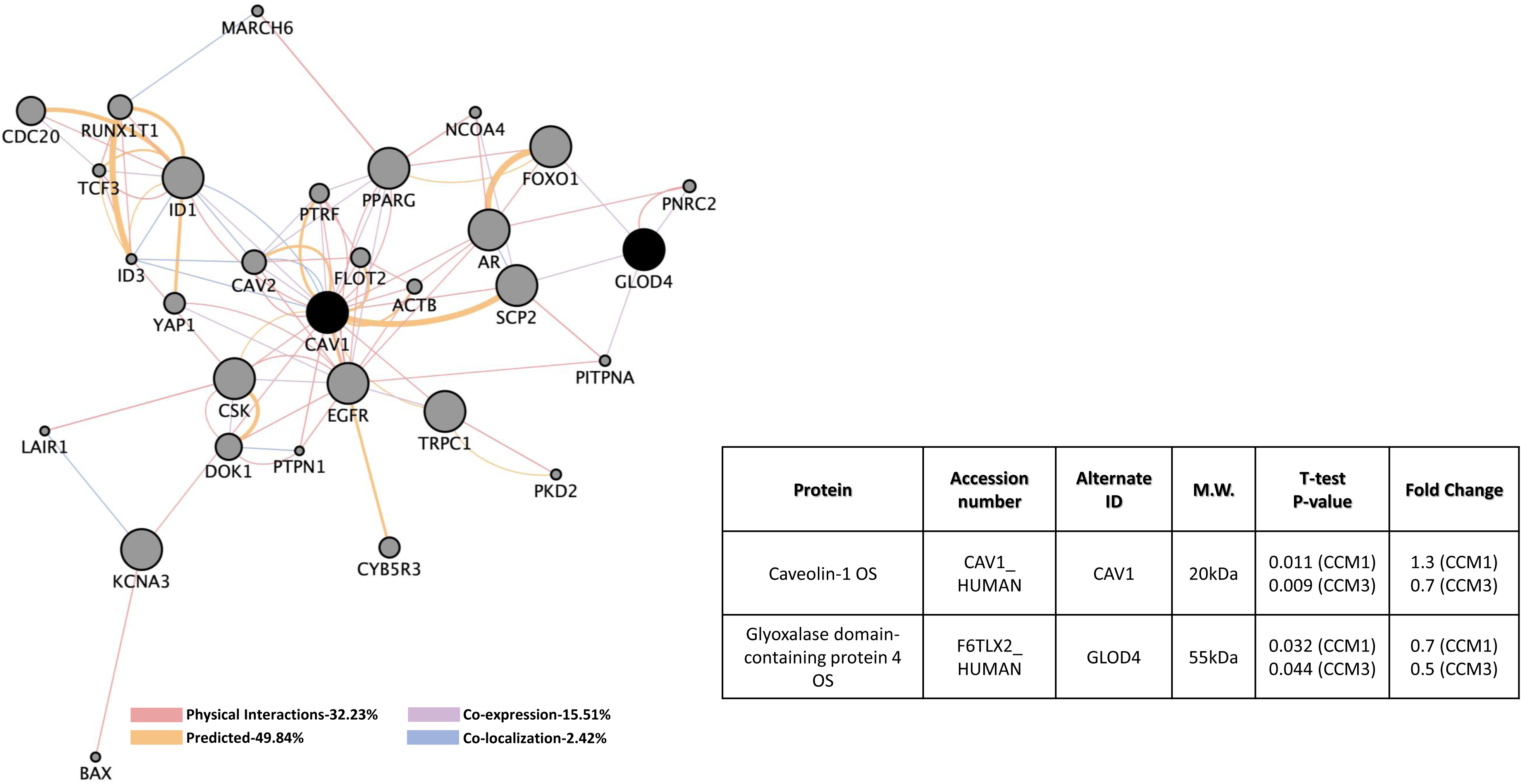
Perturbed protein interactome pathways in the deficiency of CCM1 or CCM3 in human brain microvascular endothelial cells (HBMVEC). Two proteins were identified with significant expression changes in CCM1 or CCM3 deficient HBMVEC. Table on the bottom right illustrates details of query proteins identified in the samples through mass spectrometry with accession number, alternate ID, Molecular Weight (M.W.), p-values and fold changes in mutant/controls. Interactome was built using proteomic data and constructed using cytoscape software equipped with genemania application. Interactome illustrates affected pathways with both query (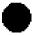) and interactors (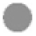). Interaction descriptions with percentages and coordinating line colors are displayed below figure. All samples were run in triplicates. Significances were determined using student’s t-test with p values <0.05.

### Perturbed protein interactome pathways shared in the deficiency of CCM2 and CCM3

Proteasome Activator Subunit 1 (PSME1), which controls cellular protein stability and found at high concentrations throughout eukaryotic cells, was down-regulated in both CCM2 and CCM3 mutants (Figure 10). Potentially affected cellular processes included various cell cycle pathways, apoptotic processes, several DNA damage pathways, antigen processing and presentation, Wnt signaling pathways, cellular response to stress pathways and several signal transduction pathways (Suppl. Table 10A). Endopeptidase activator activity functions are also associated with altered PSME1 levels (Suppl. Table 10B) as well as proteasome and antigen processing and presentation pathways (Suppl. Table 10C).

**Figure 10:**
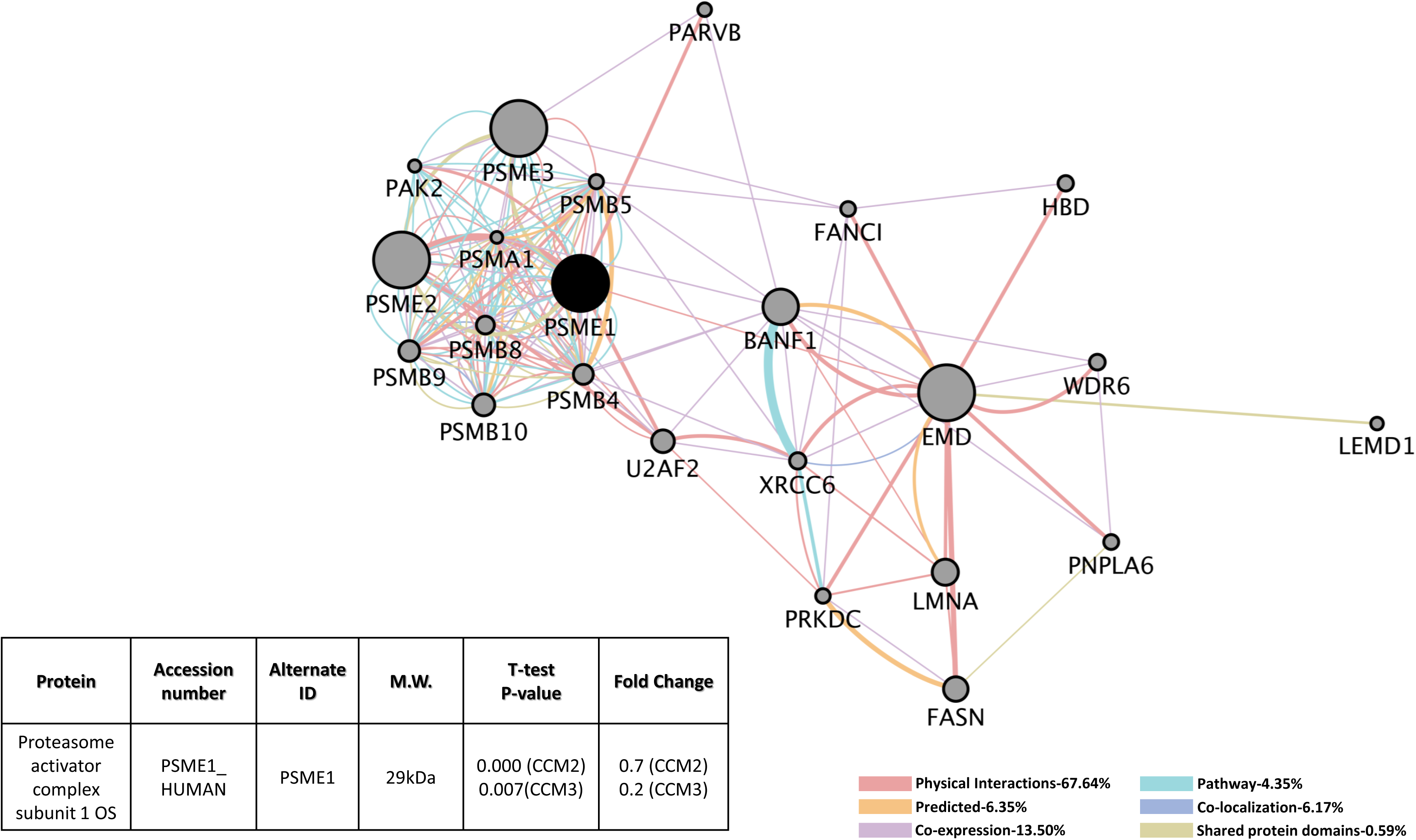
Perturbed protein interactome pathways in the deficiency of CCM2 or CCM3 in human brain microvascular endothelial cells (HBMVEC). PSME1 was identified with significant expression changes in CCM2 or CCM3 deficient HBMVEC. Table on the bottom left illustrates details of query proteins identified in the samples through mass spectrometry with accession number, alternate ID, Molecular Weight (M.W.), p-values and fold changes in mutant/controls. Interactome was built using proteomic data and constructed using cytoscape software equipped with genemania application. Interactome illustrates affected pathways with both query (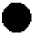) and interactors (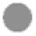). Interaction descriptions with percentages and coordinating line colors are displayed below figure. All samples were run in triplicates. Significances were determined using student’s t-test with p values <0.05.

### Perturbed protein interactome pathways shared in the deficiency of CCM1, CCM2 and CCM3

Up-regulation of SUN Domain-Containing Protein 1 (SUN1), a component of the linker of nucleoskeleton and cytoskeleton (LINC) complex that connects the cytoskeleton and nuclear lamina, was observed in all three CCM-KD HBMVEC cells (Figure 11). Interestingly, up-regulation of Glucosidase 2 (PRKCSH) in both CCM1-KD and CCM2-KD HBMVEC cells but down-regulation in CCM3-KD cells were observed (Figure 11). Lastly, DNA damage binding protein (DDB1) was also up-regulated in both CCM1-KD and CCM2-KD HBMVEC cells, but down-regulated in CCM3-KD cells (Figure 11). Numerous metabolic processes, protein modification processes, proteolysis, DNA damage response and repair, cytoskeletal anchoring at nuclear membrane, translation synthesis and cell cycle regulation pathways are potentially affected in these CCMs-KD HBMVEC cells (Suppl. Table 11A). KEGG pathways include nucleotide excision repair and ubiquitin mediated proteolysis with DDB1 altered levels in our mutant strains (Suppl. Table 11B).

**Figure 11:**
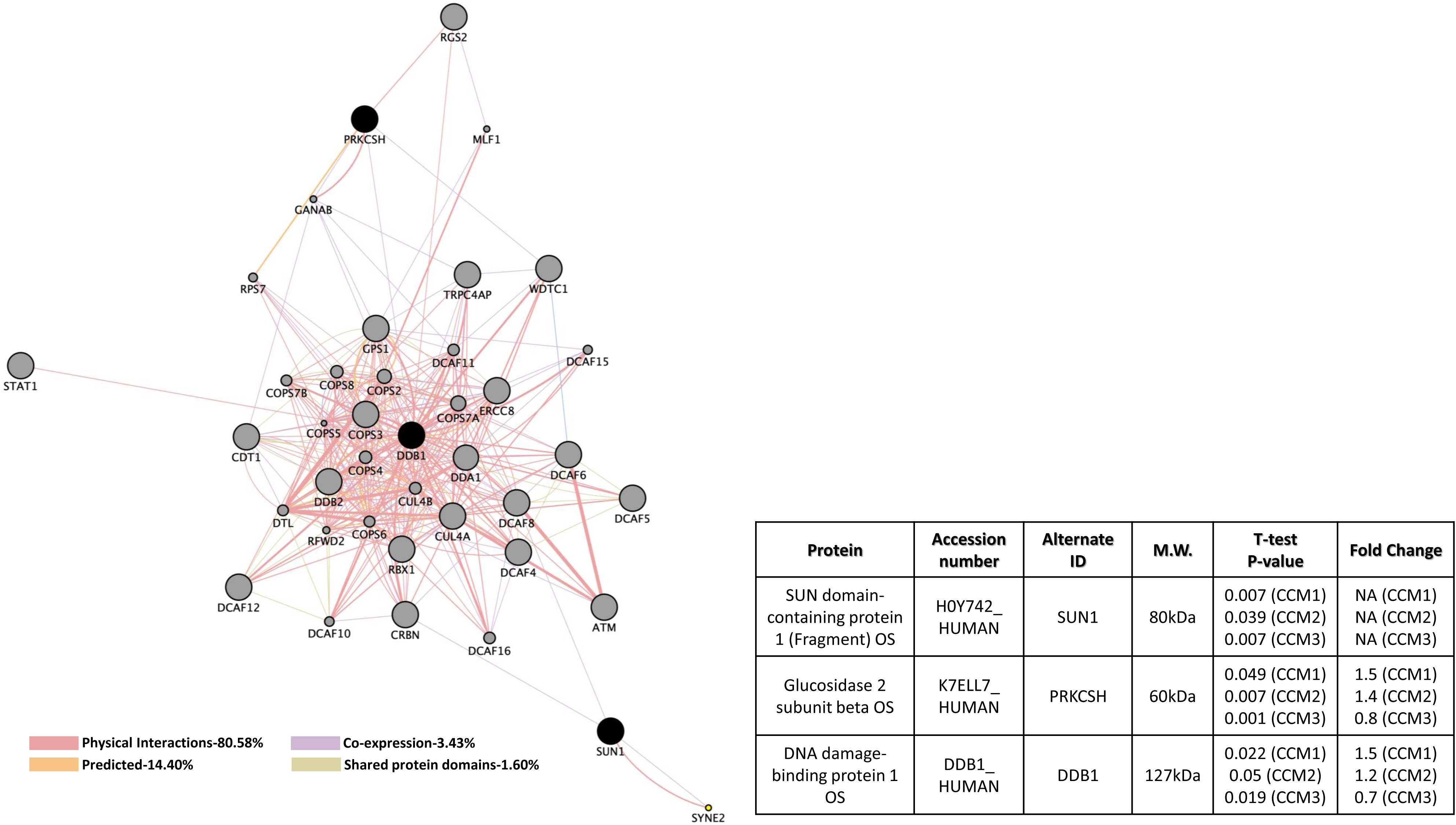
Perturbed protein interactome pathways in the deficiency of CCM1, CCM2 or CCM3 in human brain microvascular endothelial cells (HBMVEC). Three proteins were identified with significant expression changes in CCM1, CCM2 or CCM3 deficient HBMVEC. Table on the bottom right illustrates details of query proteins identified in the samples through mass spectrometry with accession number, alternate ID, Molecular Weight (M.W.), p-values and fold changes in mutant/controls. NA= not found in WT samples. Interactome was built using proteomic data and constructed using cytoscape software equipped with genemania application. Interactome illustrates affected pathways with both query (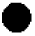) and interactors (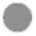). Interaction descriptions with percentages and coordinating line colors are displayed below figure. All samples were run in triplicates. Significances were determined using student’s t-test with p values <0.05.

## Discussion

### No apparent correlations between proteomic and transcriptomic results

It is highly anticipated the integrated transcriptomic and proteomic approach employed in the study will provide a systems-level understanding of cellular functions of the CSC. By comparing both transcriptomic and proteomic data, we found no overlaps between proteomic and RNA-seq results, which is not totally surprising. Messenger RNA levels do not necessarily correlate with their coded protein levels at steady state. There are several possible explanations for this discrepancy. First, there may be various regulation mechanisms existing at both RNA and protein levels for the CSC. Second, RNA-seq transcriptome profiling is not able to detect the expressional adjustment by post-transcriptional mechanisms ^64-66^. Third, there are major post-transcriptional ^67,68^ and post-translational mechanisms to separate the RNA expression and protein abundancy at static state ^69^, such as protein synthesis control pathways and protein degradation control pathways. Fourth, these results might also indicate a form of feedback inhibitory mechanisms on transcriptional regulation based on protein abundancy at static state. Finally, RNA-seq approach is more sensitive than proteomic techniques, especially in detecting low-expressed species, such as CCMs (only detected in RNA-seq in this study). However, among identified targets, there are some possible noise-like responses at both transcriptional and translational levels, due to some adaptive responses to physiological and pathological situations ^70^, unrelated to CSC alterations.

Although we were unable to find direct overlaps of altered gene expression between the RNA and protein levels, in-direct overlapped alterations were frequently seen. For example, protein level of Calpain-2 (CAPN2) was down-regulated while down-regulated RNA transcripts of Moesin (MSN), one of the interactors of CAPN2, were detected in CCM1-KD HBMVEC cells. Also, detected protein levels of Minichromosome Maintenance Complex Component 3 (MCM3) was down-regulated in both CCM1-KD and CCM2-KD HBMVEC cells while down-regulated RNA transcripts of C-C Motif Chemokine Ligand 2 (CCL2), one of the interactors of MCM3, was detected in CCM1-KD HBMVEC cells. Interestingly, in CCM2 deficient HBMVEC cells, Splicing Factor Proline and Glutamine Rich (SFPQ) protein levels were up-regulated, while down-regulated transcript levels of U2 Small Nuclear RNA Auxiliary Factor 1 (U2AF1), one of the interactors of SFPQ, was observed, suggesting that there may be a feedback regulation between SFPQ protein level and U2AF1 RNA transcript level. Our data further support the existence of multiple layers of regulation between the proteins and RNAs associated with the functions of the CSC during angiogenesis.

### Most alterations occur in angiogenic signaling cascades in perturbed CSC conditions

#### Alterations in angiogenesis at the transcript level

A significant amount of alterations at both RNA and protein levels represent an interesting outcome on potential effects concerning angiogenesis in both HBMVEC (cerebral neovascular angiogenesis) and zebrafish models (developmental angiogenesis) (Figure 12). NOV is known to play an important role in fibrosis, skeletal development and cardiovascular development ^71-76^. Several integrins (ITGαV, ITGβ3, ITGα5 and ITGβ1) are known ligands that interact with NOV to stimulate pro-antigenic activities and induce angiogenesis in endothelial cells ^77-79^. NOV RNA being down-regulated in CCM1-KD HBMVEC might suggests that the loss of CCM1 leads to reduced NOV transcription during angiogenesis.

**Figure 12:**
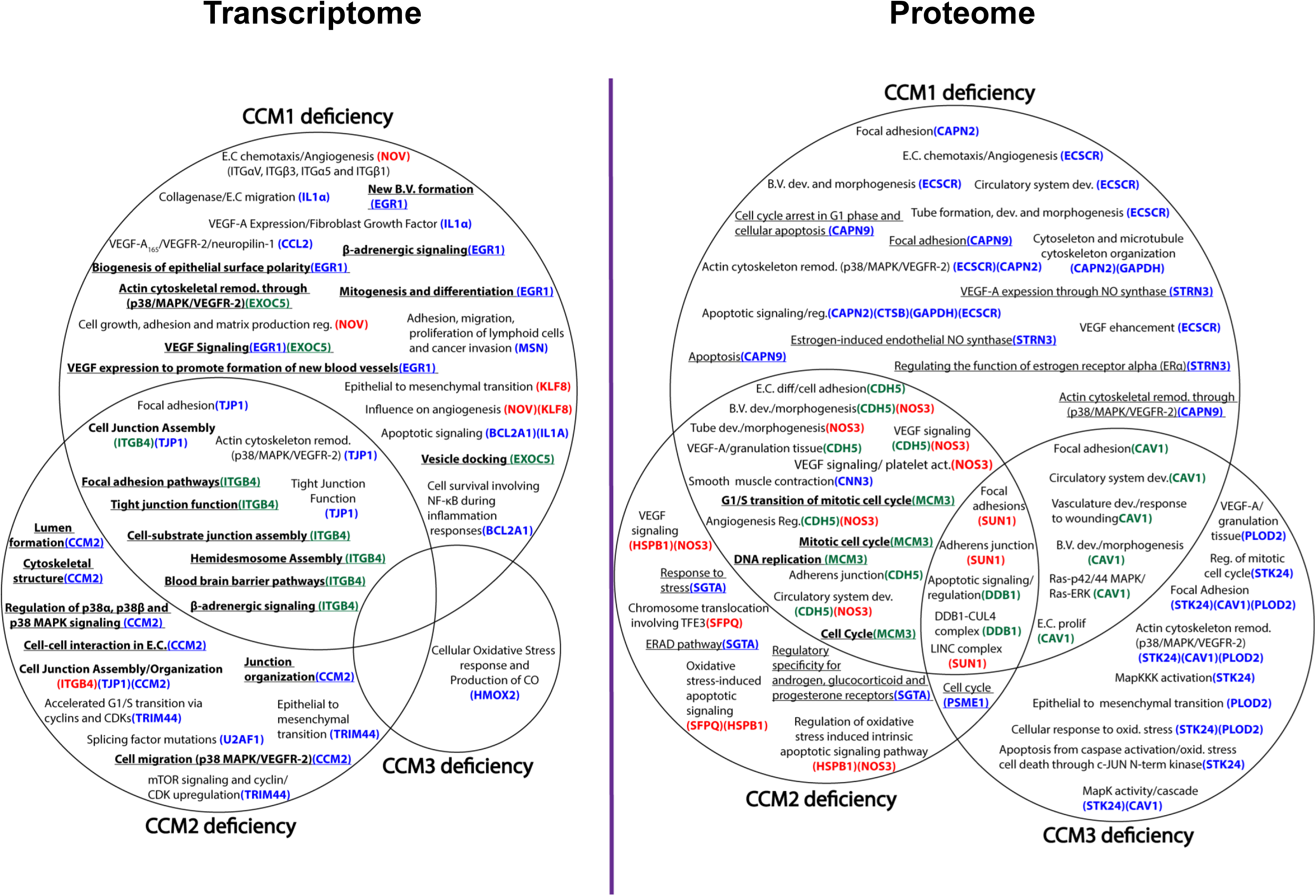
Pathways summary for angiogenesis impacts associated with perturbed CSC. Angiogenesis signaling cascades impacted with perturbed CSC in human brain microvascular endothelial cells (HBMVEC) and zebrafish strains (*san, vtn*). Venn diagram illustrates affected pathways with observed down-regulated (blue), up-regulated (red), or oppositely regulated (green) genes between CCM mutants. Diagram is divided into transcriptome (altered transcriptional data analysis) and proteome (altered translational data analysis). Pathways that are shared by more than one altered gene/protein expression is indicated with multiple genes next to that pathway. Pathways that are underlined are zebrafish specific while pathways that are in bold and underlined are shared between zebrafish (*in-vivo)* and HBMVEC (*in-vitro)* models. Abbreviations: Endothelial cells (E.C), development (dev), blood vessel (B.V.), remodeling (remod.), differentiation (diff.), activation (act.), regulation (reg.), vascular endothelial growth factor (VEGF), linker of nucleoskeleton and cytoskeleton (LINC), proliferation (prolif.) and oxidative (oxid.).

Down regulation of IL1α transcripts in CCM1 deficient HBMVEC cells potentially alters Vascular Endothelial Growth Factor A (VEGF-A) since IL1α stimulates VEGF-A expression. VEGF-A is one of the 6 members of the VEGF family which has been shown to be one of the most important factors during angiogenesis ^24^. IL1α proteins stimulate release of collagenase, a protease used to enable endothelial cell migration by breaking up the basement membrane ^36,37^. IL1α has also been shown to induce release of fibroblast growth factor (FGF), an angiogenic molecule, influencing tissue and urokinase type plasminogen activators in endothelial cells used to generate plasmin to degrade the extracellular matrix ^24,80^. FGF has also been shown to increase hypoxia-inducible protein complex HIF-1α which in turn, can promote VEGF-A expression, since this complex binds to the enhancer of VEGF-A ^24^. Alterations in both EGR1 and EXOC5 also confirmed potential effects on angiogenesis through VEGF signaling in CCM1 deficient zebrafish embryos. Focal cerebral Ischemia, one of the major diseases associated with EGR1, stimulates VEGF expression in the brain, in an effort to promote formation of new blood vessels ^23,81^. EXOC5, part of the exocyst complex, is involved in docking of vesicles on the plasma membrane ^82,83^. EXOC5 is reported to be an essential component for biogenesis of epithelial cell surface polarity, reported to interact with vesicle transport machinery and involved in actin cytoskeletal remodeling ^84^. Altered EXOC5 expression, involved in actin cytoskeletal remodeling, could also indirectly affect VEGF, since the process of cytoskeletal remodeling involves interactions of p38 MAPK with VEGFR-2 ^24^. Down-regulated CCL2, which is linked to Rheumatoid Arthritis (RA), Psoriasis and amyotrophic lateral sclerosis (ALS), is also associated with VEGF and angiogenesis signaling pathways ^24,85^. VEGF-A isoform 165, VEGFR-2 and neuropilin-1 are associated with synovial angiogenesis in RA; VEGF-A in RA will eventually stimulate generation of plasmin which leads to production of matrix metalloproteinases (MMPs) that are involved in destruction of arthritic joints characteristic in RA ^86^. VEGF-A, inhibiting apoptosis of endothelial cells (ECs) through VEGFR-2 activation, is essential in remodeling new vasculature in RA ^24^. Psoriasis is characterized by skin plaques along with neovascularization and proliferation of blood vessels; VEGF/VEGFR-1/VEGFR-2 are all overexpressed by keratinocytes in epidermis and fibroblasts influencing neovascularization at a dermal level ^24^. Both VEGF-A and CCL2 were found to be up regulated in cerebral spinal fluid in patients suffering from ALS, suggesting angiogenic responses as well as involvement of adult neural stem cells and microglial activation in pathogenesis of ALS ^85^.

Down regulation of TJP1, essential for tight junction function by coordinating binding of cytosolic proteins, F-actin and transmembrane proteins as well as a component of both human and rat BBB, was demonstrated in both CCM1 and CCM2 deficient HBMVEC. Potential alterations to focal adhesion pathways identified in CCM1 and CCM2 deficient HBMVEC interactome data, suggest a possible connection with VEGF induced cytoskeletal reorganization through its interaction of focal adhesion kinase (FAK) with paxilin ^87^. Altered protein expression of ITGB4 was observed in CCM1 and CCM2 deficient HBMVEC as well as CCM1 deficient zebrafish, indicating an important role of ITGB4 in the CSC. ITGB4 signaling pathway plays an essential role in the integrity of the BBB. This data suggests overlaps in potential alterations in tight junction, focal adhesion and BBB pathways when either CCM1 or CCM2 expression levels are altered in HBMVEC and zebrafish model.

#### Potential alterations in angiogenesis at the proteome level

Correlating with affected pathways concerning angiogenesis at the transcription level, there was a number of affected pathways at the protein level that reinforces CCM genes’ involvement in angiogenesis in both HBMVEC and zebrafish models (Figure 12). Down-regulated in CCM1-KD, ECSCR is known to regulate tube formation as well as endothelial chemotaxis and interact with filamin A ^88^, and is involved in EGF-induced cell migration and cell shape alterations. ECSCR is primarily found in blood vessels and ECs ^89^, asserting its role in angiogenesis through modulation of actin cytoskeleton, and enhances VEGF ^90^. Down-regulation of CAPN2 potentially affects focal adhesion pathways associated with CAPN2. This data, similar to that mentioned above with RNA transcript analysis of CCM1 and CCM2 deficient models, suggests a possible connection with VEGF induced cytoskeletal reorganization ^87^. Down**-**regulation of CAPN2 and GAPDH illustrated potential alterations to regulation of cytoskeleton and microtubule organization. Down-regulated Striatin 3 (STRN3), unique in zebrafish Ccm1 mutant only, is suspected to function as a signaling or scaffold protein, and shown to have a role in regulating the function of estrogen receptor alpha (ERα) and acts as a scaffold for assembly of proteins necessary for rapid activation of ERα-induced eNOS ^91^. VEGFR-2 also activates eNOS which aids in generating nitric oxide to increase vascular permeability and cellular migration. Overexpression of cyclooxygenase (COX)-2, an enzyme induced by eNOS, stimulates tube formation and EC motility through increased production of VEGF-A ^24^. Up-regulation of Heat Shock Protein Family B Small Member 1 (HSPB1), as seen in CCM2 deficient HBMVEC lines, is shown to be involved in VEGF signaling pathways. Up-regulation of eNOS, in both CCM1 and CCM2 HBMVEC mutants, promotes blood clotting by activating platelets, and participating in VEGF induced angiogenesis ^54^. Further demonstration of alterations in angiogenesis involving eNOS include potentially affected pathways involved in regulation of blood circulation, blood vessel diameter, blood coagulation, blood vessel EC migration, and platelet activation. VE-Cadherin, up-regulated in CCM1 and down-regulated in CCM2 deficient HBMVEC, is a key player in endothelial adherens junction assembly and maintenance. VEGF-A disorganizes endothelial junction proteins, including cadherin proteins such as VE-Cadherin, to allow for decreased barrier properties and gap formation between ECs, thereby enhancing supply of proteins and cells to form granulation tissue ^24^. VE-Cadherin alterations potentially affect pathways including cell-cell adhesion and regulation of EC differentiation. Altered pathways involving both eNOS and VE-Cadherin expression levels include blood vessel development, circulatory system development, VEGF receptor signaling pathway and regulation of angiogenesis. Calponin-3 (CNN3), down-regulated in both CCM1 and CCM2 HBMVEC mutants, is a cytoskeleton protein and has been shown to regulate actin cytoskeleton functions and smooth muscle contraction. When looking at CCM3 deficiency, down-regulation of STK24 expression levels present a potential effect on axon regeneration in the radial and optic nerves. STK24 promotes apoptosis in response to caspase activation and stress stimuli, as well as mediating oxidative stress induced cell death by modulating c-Jun N-terminal kinase (JNK) activation ^92,93^. In regards to angiogenesis, binding of VEGF to VEGFR-2 results in a downward signaling cascade that eventually activates p42/44 MAPK that phosphorylates and activates c-JUN ^24^. Altered gene expression of STK24 affects several kinase cascade pathways including activation of MAPKKK and various regulation of both MAPK activity and MAPK cascade. VEGF-A and its target receptors, VEGFR-1 and VEGFR-2, undergoes increased expression when cells are undergoing hypoxia; in several *in-vivo* models ^24^. Interestingly, alterations in STK24 affect several pathways associated with oxidative stress including intrinsic apoptotic signaling pathways in response to oxidative stress.

Down-regulation of PLOD2 is also expected to alter angiogenesis through VEGF induced cytoskeletal reorganization through VEGFR-2/p38MAPK/focal adhesion interactions in HBMVEC lines. PLOD2 facilitates hydroxylation of lysyl residues in collagen-like peptides and is involved in degradation of the ECM and collagen chain trimerization, an important step for transforming fibrin-fibronectin stroma into granulation tissue of healing wounds ^24^. PLOD2 is also implicated in response to oxidative stress, similar to STK24 mentioned above. One of the main functions of the multidomain scaffolding protein TJP1, is to help regulate movement of macromolecules and ions between endothelial and epithelial cells ^94^. TJP1is essential for tight junction function, and has been shown to be a key component for human and rat BBB integrity ^95-97^.

Altered protein expression of CAV1 was defined in CCM1 and CCM3 deficient cells, CAV1 negatively regulate Ras-p42/44 mitogen-activated protein kinase (MAPK) cascade, promotes cell cycle progression and participates in coupling integrins signaling to the Ras-ERK pathway. The p42/44 MAPK pathway is involved as a downstream effect to the binding of VEGF to VEGFR-2 and also to binding of VEGF to VEGFR-3. Altered levels of CAV1 could therefore indirectly provide some feedback inhibition signaling for altered VEGF/VEGFR-2 binding, influencing blood vessel EC proliferation, migration and survival; while VEGF/VEGFR-3 binding, influences survival of blood and lymphatic EC survival, migration and proliferation ^24^. Additionally, multiple pathways potentially affected by CAV1 expression, reinforces its role in angiogenesis, including circulatory system development, blood vessel development and morphogenesis, vasculature development, nitric oxide biosynthetic process, response to wounding, focal adhesion pathways and EC proliferation.

Shared alterations in SUN1 levels were observed in CCM1, CCM2 and CCM3 deficient HBMVEC cells, suggesting its major role in CSC-mediated angiogenesis signaling. SUN1 is a component of the linker of nucleoskeleton and cytoskeleton (LINC) complex that connects the cytoskeleton and nuclear lamina. The cytoskeleton, important for various steps in angiogenesis, cell motility and membrane protrusion events are essential for establishing successful angiogenic responses ^98^. Furthermore, it has been demonstrated that the focal adhesions at cell-matrix and adherens junctions at cell-cell contacts that tether the cytoskeleton to the plasma membrane are essential for angiogenic responses; it was demonstrated that when cadherin’s, focal adhesions and integrins are inhibited, angiogenic responses are blocked ^99-101^.

### Altered β4 integrin signaling in all perturbed CSC conditions is a surprise

Perturbation of integrin β-4 (ITGB4) signaling was consistently observed in both CCMs-KD HBMVEC cells and CCMs-KO zebrafish embryos; which is the only signaling cascade validated in both human and animal models (Figures 3A and 3B). Up-regulation was observed among CCM1-KD and CCM2-KD HBMVEC cells and Ccm1-KO zebrafish (*san*) but down-regulated in Ccm2-KO zebrafish (*vtn*) (Figures 3A and 3B). ITGB4 is different from other β subunits for its extra-large cytoplasmic tail (over 1,000 amino acid long), docking to intermediate filaments through plectin ^32,102^, while cytoplasmic tails of other β subunits (∼ 50 amino acids) bind to actin cytoskeletons. As a laminin-5 (LN5) receptor and a key component of hemidesmosome (HD) ^103^, ITGB4 is mainly located in the adhesion structure of HD, contributing to the maintenance of integrity of epithelium ^102^, especially the epidermis ^33,104^. In angiogenesis, ITGB4 was initially found to be located predominantly on the matured vasculature where it is negatively regulated, suggesting ITGB4 as a negative regulator of angiogenesis at quiescent state ^105^. Up-regulated integrin β4 was suggested to be associated with vascular disease related to aging ^106^.

One mutagenesis study demonstrated that ITGB4 could promote cancer angiogenesis in tumorigenesis ^31^, suggesting its potentially versatile roles in angiogenesis in various conditions. ITGB4, which binds to the basal lamina of blood vessels, was initially found to be expressed in small vessels ^105,107^ but recently, ITGB4 was found to be highly expressed in the brain. Initial studies suggested that ITGB4 is expressed by astrocyte end-feet that run along the vascular basal lamina ^108,109^, which was supported by animal studies showing transgenic mice deficient in astrocyte or pericyte laminin displayed defective BBB integrity ^110,111^. Pathways associated with ITGB4 include BBB and β-adrenergic signaling pathways ^112,113^. A recent study found that ITGB4 is highly expressed in ECs in cerebral vessels, specifically in arteriolar ECs ^113^, raising the possible existence and important role of the adhesive HD structure in cerebrovascular ECs and BBB. However, further study found that ITGB4 is not essential for localization or regulation of expression of key HD structural proteins, plectin and CD151 ^114^, making the function of ITGB4 interestingly elusive in cerebral vessels.

Up-regulation of ITGB4 observed in CCM1 and CCM2 deficient HBMVEC as well as CCM1 deficient zebrafish, but down regulation in CCM2 deficient zebrafish strains offer an interesting and contradicting regulation mechanism for ITGB4 whose main pathways lay on cerebral vessels and the integrity of BBB. Since three CCM proteins influence each other within the CSC, there might be a “paradoxical regulation of ITGB4”, mimicking regulation of CAV1, as described below. Therefore, in the perturbed CSC, we postulate that ITGB4 likely functions as a cellular sensor guarding for the integrity of BBB in cerebrovascular ECs. In response to environmental cues, CSC signaling can modulate BBB status by regulating ITGB4 expression level in ECs. In an effort to enhance the weakened BBB integrity of cerebral vessels, ITGB4 expression in ECs can be enhanced by CSC, increasing the ability of anchoring to the basal lamina. ITGB4 also plays many other important roles in ECs cell physiology, including survival, proliferation, apoptosis, autophagy, angiogenesis, and senescence. Recently, reported up-regulation of ITGB4 expression in cerebrovascular ECs has been associated with neuroinflammatory events ^115-117^, raising the possible role of ITGB4 in connection between neuroinflammation and perturbation of CSC as discussed below.

### CSC might be a master-regulator for CAV1-mediated signaling

CAV1 is an essential component of caveolae that are 60–80 nm wide, bulb-shaped pits in the plasma membrane ^118^. Each caveola has an estimated 140–150 Cav1 molecules, which usually form oligomers in the plasma membrane ^119,120^; genetic ablation of CAV1 causes loss of caveolae ^121,122^. It has been demonstrated that β1 integrin regulates caveola formation ^123^, and caveolar density decreases when integrin-mediated adhesion to the extracellular matrix (ECM) is disrupted ^124^. Reciprocally, caveolin can also contribute to β1 integrin endocytosis to influence β1 integrin signaling in either positive ^125,126^ or negative ^127^ fashions, depending on specific cell types ^128^. Cav1 plays an important role in endocytosis, and caveolin-dependent endocytosis is modulated through RAC1– p21-activated kinase (PAK), PI3K–AKT and RAS–ERK signaling cascades ^124,129^. These reports are fully in concordance with our previous observations that perturbation of CSC signaling leads to disruption of β1 integrin-mediated angiogenesis ^11,13,19-22,130^, altered RAC1–p21-activated kinase (PAK), PI3K–AKT and RAS–ERK signaling cascades ^19,21,22,130,131^. This data strongly suggests that CAV1 signaling is downstream of CSC signaling complex.

It has been well defined that CAV1 directly interacts with endothelial nitric oxide synthase (eNOS) through its binding to caveolin-binding motifs (CBM) in eNOS ^132,133^ and inhibits eNOS signaling ^133-135^. Intriguingly in this study, eNOS was found to be up-regulated in both CCM1-KD and CCM2-KD HBMVEC cells (Figure 7A), further enhancing our previous conclusion that CAV1 signaling is modulated by CSC as one of its downstream signaling targets. Interestingly, CAV1 protein level was up-regulated in CCM1 deficiency, while down-regulation of CAV1 protein in CCM3 depletion was observed (Figure 9). This contradicting phenomena has been well described as the caveolar “paradox” ^136^. In this paradoxical regulation, there is a coexistence of the “signaling inhibitory” paradigm allosteric inhibition of eNOS proportional to caveolin levels and the “signaling promoting” effect of caveolin through compartmentation of eNOS according to the abundance of caveolin. Therefore, there is a typical bell-shaped relationship between the abundance of CAV1 and NO-driven angiogenesis, in which there is only a narrow center-region of pro-angiogenic effects for CAV1-eNOS signaling. Anti-angiogenic effects will be observed when CAV1 expression level shifts out of the normal range, as up-regulation leads to an exacerbated inhibitory clamp, while down-regulation leads to loss of compartmentation ^137^. Therefore, CSC can be a master regulator to either positively (via CCM3) or negatively (via CCM1) regulate CAV1 expression to fine-tune the intricate balance of CAV1-eNOS signaling cascades.

Likewise, canonical Wnt signaling is essential for both brain specific angiogenesis and blood brain barriergenesis ^138,139^, in which β-catenin signaling may mediate BBB functions ^140^, and phosphorylated CAV1 can sequester β-catenin and thus decrease BBB function ^141^. Interestingly, pathway analysis showed that perturbed Wnt signaling pathways were found to be shared in the deficiency of CCM2 and CCM3 (Suppl. Table 10A). One report described that EC-specific Ccm3-KO led to activation of β-catenin, preceding the initiation of TGF-β/BMP signaling ^142^, which might add more evidence that CSC signaling is upstream of Wnt signaling pathways in modulating the activity of β-catenin. These observations have expanded our view about the relationship and signal flow direction among CSC signaling, CAV1 signaling and Wnt signaling cascades during angiogenesis.

Oxidative stress has been defined as molecular insults resulting from excessive and/or cumulative production of reactive species, such as reactive oxygen species (ROS) and reactive nitrogen species (RNS) ^143^. Cav-1 has been shown to have regulatory feedback functioning on oxidative stress status. As discussed before, as a molecular hub, Cav-1 can bind and negatively regulate the activity of eNOS, therefore, loss of Cav-1 will lead to up-regulation of eNOS activity and increased NO production, which could lead to accumulation of RNS. Further, Cav-1 regulates a variety of cellular signaling, such as RAs-MAPKs signaling ^144^ and AKT signaling pathways ^145,146^, therefore silencing Cav-1 could result in enhanced oxidative stress ^147^.

Chronic inflammation has been proven to be associated with elevated oxidative stress in cardiovascular diseases ^148^. A group of inflammatory factors, including CCL2, IL1α, MSN, BCL2A1 and LMAN1, were all significantly down-regulated in CCM1-KD HBMVEC cells. CCL2 has a specific role as a chemotactic factor attracting basophils and monocytes ^149^. CCL2 expression is up-regulated during degenerative and inflammatory conditions in the CNS ^150^, and is associated with RA, atherosclerosis and psoriasis ^151,152^. IL1α, through maturation and proliferation of B-cells and induction of IL-2 release and fibroblast growth factor activity, is able to stimulate inflammation ^80,153,154^. Being involved in inflammatory responses, IL1α has been reported to stimulate release of collagenase and prostaglandin from synovial cells ^36,37^. MSN, involved in adhesion, migration and proliferation of human lymphoid cells, are known to contribute to immunologic synapse formation ^155-157^. RhoA signaling and actin nucleation by ARP-WASP are all pathways involving participation of MSN ^158,159^. Dysregulation of MSN might be associated with immunodeficiency ^160,161^. BCL2A1, being a direct target of NF-κB during inflammation responses, is believed to have a cyto-protective function essential for cell survival and lymphocyte activation; BCL2A1 is up-regulated in the presence of signals such as IL1α and TNF ^162-164^.

Previous reports indicate that CCM1 plays a protective role against oxidative stress in the cell by utilizing anti-inflammatory and anti-oxidant pathways ^165^. CCM1 deficient cells have increased activity of COX-2, a mediator of inflammatory pathways ^166^; furthermore, Ccm1-KO mice had hyper-exaggerated responses to inflammatory agents ^167,168^, suggesting that neuroinflammatory events might be associated with CCM lesion formation ^169^. Further, we observed that regulation of oxidative stress-induced signaling pathway was changed in CCM2 deficient HBMVEC. This current study falls in line with these previous findings.

### Protein stability regulation in perturbed CSC condition

Heat shock proteins (HSPs), mainly functioning as chaperones, whose altered expression levels can result in misfolded proteins and lead to degradation. Up-regulation of both HSPA4 and HSPB1 was observed in CCM2 deficient HBMVEC lines, contradicting to previous report that CCM1 deficient cells have decreased levels of heat shock proteins (HSP, Hsp70 and Hsp27), which have a protective role in handling cellular stress ^170^. DDB1, perturbed in all three CCMs-KD HBMVEC cells, is a part of the DDB1-CUL4 ubiquitin ligase complex, a key factor in autophagy. Autophagy removes protein aggregates and damaged organelles from cells and helps to recycle lipids and protein building blocks, a constitutively active, evolutionary conserved, catabolic process for maintaining homeostasis in cellular stress responses and cell survival. Regulatory mechanisms of this process are still largely unknown. LMAN1, down-regulated in CCM1-KD HBMVEC cells, functions as a cargo receptor for glycoprotein transport, cycles between cis-Golgi, the endoplasmic reticulum (ER), and the ER-Golgi intermediate compartment ^171^, which might participate in ER stress-induced autophagy. SGTA, down-regulated in CCM2-KO zebrafish, is not characterized very well but has been shown to interact with 70-kDa heat shock cognate protein and hypothesized to serve housekeeping functions ^172,173^. DHRSX, up-regulated only in CCM1 deficient cells, is relatively uncharacterized other than being hypothesized in having a role in positive regulation of starvation-induced autophagy ^174^. Previous data showed that down-regulation of integrin β4 promotes sphingosylphosphorylcholine-induced autophagy ^175^, suggesting that perturbed autophagy can be observed in CCMs. Therefore, this data is in agreement with previous reports that defective autophagy is a key feature of CCMs ^176^.

### Endothelial-to-mesenchymal transition factors in perturbed CSC

Originally found in tumor research, endothelial-to-mesenchymal transition (EndMT) is a phenomena in which ECs change its organized vascular EC cell layer phenotype in losing cell–cell junctions into a mesenchymal phenotype of invasive and migratory properties to invade the underlying tumor tissue and is one of the major causes of fibro-proliferative vascular disease ^177-180^. EndMT accounts for up to 40% of cancer-associated fibroblasts (CAFs). EndMT is often considered as a specialized form of epithelial-to-mesenchymal transition (EMT), which has been extensively studied in cancer. Both EMT and EndMT generate cells that share a similar mesenchymal phenotype and utilize common signaling pathways ^181^. Therefore, both share the same, if not identical, mesenchymal markers such as fibroblast-specific protein 1 (FSP1/S100A4), α-smooth muscle actin (αSMA) or a group of transcription factors (Snail, Zeb1/2, Twist, Slug), making it difficult to distinguish both events ^182,183^.

High PLOD2 expression is associated with EMT ^184,185^. Likewise, TRIM44 acts as an EMT-promoting molecule ^186,187^, with overexpression of TRIM44 inducing EMT transition ^188,189^. It has been reported that perturbed CSC was able to induce EndMT in brain microvascular ECs that plays a crucial role in the pathogenesis of CCMs ^190^. It was further reported that EC-specific Ccm3-KO led to activation of β-catenin preceding the initiation of TGF-β/BMP signaling. A β-catenin-driven transcription program that leads ECs to EndMT in CCM3 deficient ECs, and pharmacological inhibitors of β-catenin signaling can significantly reduce progress of this EndMT transition ^142^. Then, it was reported that transcription factors KLF 2/4 (Kruppel-like factor 2 or 4) can be strongly induced in CCM1-null ECs, which, in turn, promotes TGFβ/BMP signaling, leading to EndMT ^191-194^. These findings support a theory that the CSC has an inhibitory effect on the EndMT/EMT cellular processes.

Surprisingly in our data, the significantly changed expression levels of two EndMT/EMT factors identified, PLOD2 in CCM3-deficinet HBMVEC and TRIM44 in CCM2-deficinet HBMVEC cells, were both down-regulated, contradicting to the previous theory. Likewise, down-regulation of both TJP1 and ITGB4 was reported in platelet-derived growth factor–D (PDGF-D) induced EMT ^195^. Interestingly, we observed down-regulation of TJP1 in both CCM1-KD and CCM2-KD HBMVEC cells and down-regulation of ITGB4 in Ccm2-KO zebrafish (*vtn*), however, up-regulation of ITGB4 was found among CCM1-KD, CCM2-KD HBMVEC cells and Ccm1-KO zebrafish (*san*) (Figures 3A and 3B), discordant to previous results. Although no altered expression of reported EndMT/EMT factors, KLF 2/4 was found in our data, increased expression of KLF8 was found in CCM1 deficient HBMVEC cells; whether this is a KLF 2/4 related pathway or a feedback regulation in the defective EndMT/EMT needs to be further investigated. In sum, our systems biological data suggests that pathogenesis of CCM seems unlikely to be directly associated with EndMT/EMT pathways in our current models.

### Neurological conditions associated with perturbed CSC

CCM itself, as a neurological disorder, can certainly be associated with many other neurological conditions. APBA2 and CTSB are known to interact with amyloid precursor protein (APP) associated with Alzheimer’s disease (AD). CTSB, down-regulated in CCM1-KD HBMVEC cells, is known for its role in proteolytic processing of APP, in which disruption of this process is suggested to play a major role in AD ^196-198^. Interestingly, APBA2 also down-regulated in CCM1-KD HBMVEC cells, stabilizes APP to stop production of proteolytic fragments of APP often found in the brain of patient’s suffering from AD ^199-201^. APBA2 is also hypothesized to function as a vesicular trafficking protein that couples synaptic vesicle exocytosis to neuronal cell adhesion ^202-204^. AD is associated with dysregulated CAPN2 gene, and this family of proteins have been shown to play a major role in neuropathological events following traumatic brain injury ^205-209^. Perturbed expression levels of HSP protein can often be observed in ALS, AD and Huntington’s disease. HSPA4 is known to interact with Parkin, PINK1 and DJ-1 of which all three genes have been linked with Parkinson disease and has been shown to interact with transcription factor C-myc, Creb1 and Egr1 ^210^. GAPDH, down-regulated in CCM1-KD HBMVEC cells, by stimulating binding of CHP1 to microtubules, plays a major role in CHP-1 dependent microtubule and membrane associations ^211^. GAPDH is also a key player in assembly and organization of the cytoskeleton, and has been implicated in AD-related apoptotic cell death ^212,213^.

eNOS, up-regulated in both CCM1 and CCM2 HBMVEC mutants, mainly functions to produce nitric oxide. AD is also associated with altered eNOS expression levels ^55-59^. PRPF8, up-regulated in CCM1-KD HBMVEC cells, has been reported to be essential in pre-mRNA splicing processes, cell survival and myeloid differentiation ^214^. This protein is associated with rare, genetic disorders involved in the breakdown and loss of cells in the retina ^215,216^. TALDO1 is known to be a key player in both nucleic acid synthesis as well as lipid biosynthesis ^217^. Interestingly, multiple sclerosis (MS), transaldolase deficiency (TALDO1) and Ribose 5-Phosphate Isomerase Deficiency (RPIA) are associated with TALDO1 ^218^.

## Conclusion

Although there have been many efforts in elucidating the molecular and cellular effects of deficiency of each CCM protein, currently, there is no report yet on dissecting the perturbed CSC by utilizing a systems biology approach. This first effort of systems analysis on CCMs deficient models in HBMVEC (*in-vitro)* and zebrafish lines (*in-vivo*), elucidates a very unique and interconnected story demonstrating alterations in various signaling cascades both in cerebral neovascular angiogenesis as well as developmental angiogenesis with disruption of the CSC. These alterations mainly involved angiogenesis signaling cascades, VEGF pathways, β4 integrin signaling, CAV1 signaling, autophagy and association with altered inflammatory factors (Figures 12 and 13). Alterations in angiogenic pathways mainly involved in 1). *Angiogenic cues*: endothelial cell chemotaxis, ECM, VEGF signaling, FGF signaling; 2). *Blood vessel formation*: cytoskeletal remodeling, endothelial cell migration, lumen formation, tube morphogenesis and development, vessel morphogenesis and development; 3). *blood vessel maintenances*: Blood circulation, regulation of blood vessel diameter, blood coagulation; 4). *Cell junction organization*: Focal adhesion pathways and tight junction function; and 5). *P38/MAPK cascade, response to oxidative stress* (Figure 12 and 13). To further elucidate these altered pathways, *in-vivo* mouse models in the deficiency of CCMs will be utilized to evaluate the impact of alterations of the CSC with various approaches to further dissect the underlying mechanism of CSC signaling during angiogenesis.

**Figure 13:**
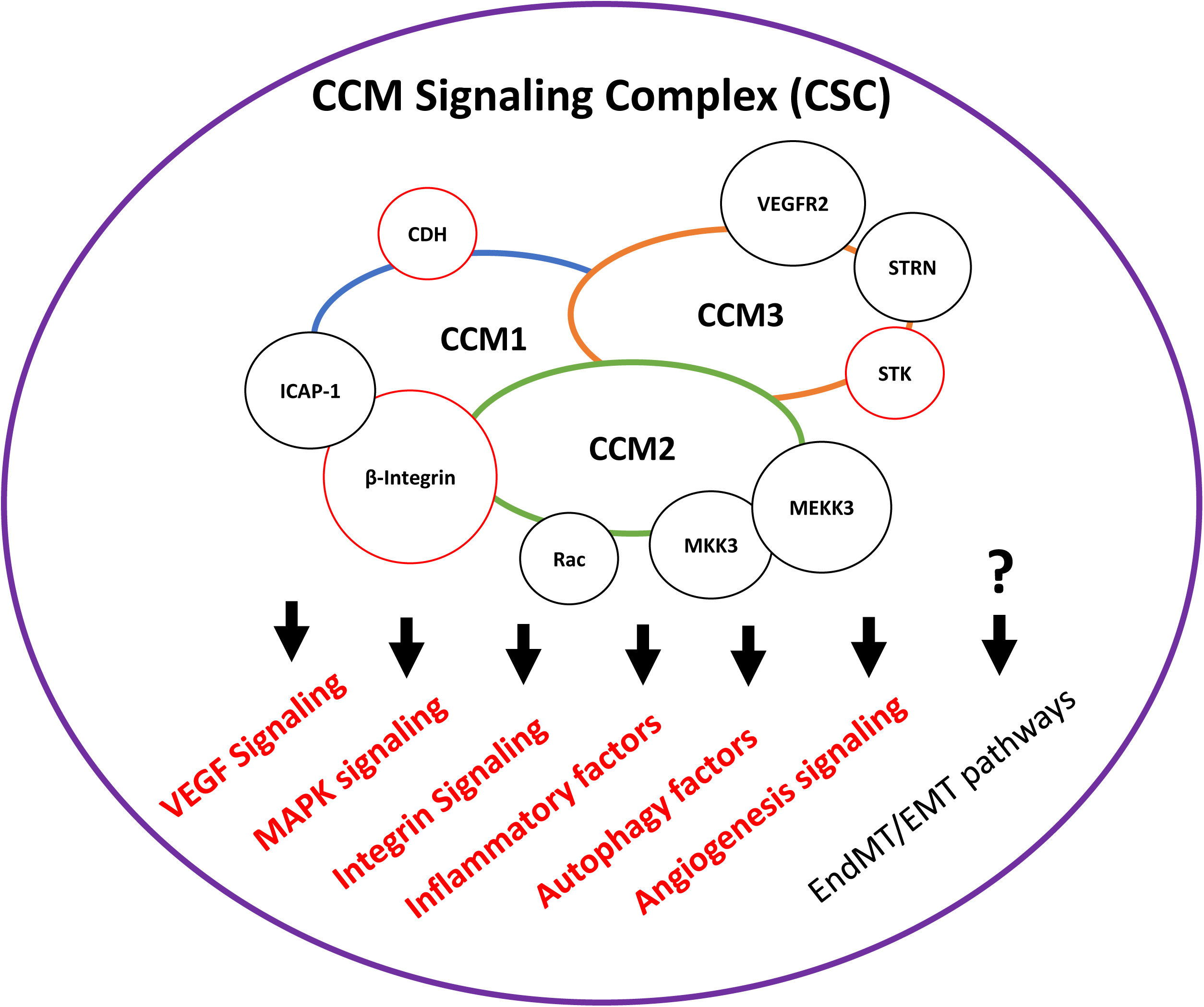
Summarized major signaling pathways associated with perturbed CSC. Pathways impacted in our models with perturbed CSC signaling in human brain microvascular endothelial cells (HBMVEC) and zebrafish strains (*san, vtn*) in this study are labeled in red beneath the diagram. Diagram illustrates affected pathways in regards to angiogenesis with interactors bound to each CCM protein; interactors in red were found at altered levels in our current CCM mutagenesis study. Signaling cascade not supported in this study is labeled with a question mark.

## Abbreviations

CCM: Cerebral cavernous malformation
CSC: CCM signaling complex
HBMVEC: human brain microvascular endothelial cells
VEGF: zebrafish Ccm1 (*san*) and Ccm2 (*vtn*) mutant strains, vascular endothelial growth factor
MAPK: mitogen activated protein kinase
CNS: central nervous system
ECM: extracellular matrix
BBB: blood brain barrier
EC: endothelial cell

## Acknowledgments

We wish to thank Junli Zhang, Nancy (Xiaoting) Jiang, Akhil Padarti, Yanchun Qu, Shen Sheng, Ahmed Badr, and Elias Gonzalez at Texas Tech University Health Science Center El Paso (TTUHSC) for their technical help during the experiments.

## Author Contributions Statement

J.A.F and J.Z contributed in writing the main manuscript text. J.Z, and J.A.F contributed in experimental and material generation and preparations for genomic and proteomic studies. M.V. and J.A.F contributed in RNA-seq analysis while B.G, C.E and J.A.F contributed in proteomic assay and data analysis. J.A.F contributed in integrated systems biology analysis. J.Z contributed in overall experiment design and execution. All authors reviewed the manuscript.

## Compliance with Ethical Standards

### Ethical approval

This article does not contain any studies with human participants performed by any of the authors, nor studies with animals performed by any of the authors.

